# Multifactorial heterogeneity of the human mutation landscape related to DNA replication dynamics

**DOI:** 10.1101/2022.09.28.509938

**Authors:** Madison Caballero, Dominik Boos, Amnon Koren

## Abstract

Mutations do not occur uniformly across genomes but instead show biased associations with various genomic features, most notably late replication timing. However, it remains contested which mutation types in human cells relate to DNA replication dynamics and to what extents. Previous studies have been limited by the absence of cell-type-specific replication timing profiles and lack of consideration of inter-individual variation. To overcome these limitations, we performed high-resolution comparisons of mutational landscapes between and within lymphoblastoid cell lines from 1662 individuals, 151 chronic lymphocytic leukemia patients, and three colon adenocarcinoma cell lines including two with mismatch repair deficiency. Using cell type-matched replication timing profiles, we demonstrate how mutational pathways can exhibit heterogeneous replication timing associations. We further identified global mutation load as a novel, pervasive determinant of mutational landscape heterogeneity across individuals. Specifically, elevated mutation load corresponded to increased late replication timing bias as well as replicative strand asymmetries of clock-like mutations and off-target somatic hypermutation. The association of somatic hypermutation with DNA replication timing was further influenced by mutational clustering. Considering these multivariate factors, and by incorporating mutation phasing at an unprecedented scale, we identified a unique mutational landscape on the inactive X-chromosome. Overall, we report underappreciated complexity of mutational pathways and their relationship to replication timing and identify specific factors underlying differential mutation landscapes among cell types and individuals.

## Introduction

Mutations arise through a compendium of known and unknown mechanisms. These include the improper repair of DNA damage produced by endogenous or exogenous agents, enzymatic alterations of DNA, and mismatches introduced during DNA replication. Knowing how, where, and when mutations occur is central to understanding evolution, aging, and disease. In this respect, it is well established that mutations are distributed non-randomly at the nucleotide, regional, and global genomic levels. At the nucleotide level, many mutational pathways are biased toward specific nucleotide substitutions and surrounding sequence contexts^1^. For example, the spontaneous deamination of 5-methylcytosine to thymine happens almost exclusively at CpG sites^2^. On a regional and global scale, variations in mutation rates and substitution types are associated with various genetic and epigenetic factors including nucleotide content^3,4^, chromatin state^5–7^, three-dimensional genome organization^8^, transcription factor binding^9,10^, and DNA replication timing^11–20^.

DNA replication timing is the cell type-specific spatiotemporal pattern of genome replication along S-phase. In eukaryotic cells, DNA replication begins at multiple replication origins that fire throughout S-phase and mediate bidirectional replication until the entire genome is duplicated. Late replicating regions of the genome are broadly enriched for single nucleotide variants and mutations^11,12,14–16,21,22^. The mechanisms by which mutations accumulate in later replicating regions of the genome remain incompletely understood, although evidence suggests that mismatch repair (MMR) attenuates toward the end of S-phase and contributes to these biases ^16,23^. On the other hand, many classes of mutations and their underlying mutational pathways are not biased with respect to replication timing^12,15^, suggesting complex contributions by different DNA damage and repair pathways.

A powerful method to glean the types and abundances of mutational pathways that shape mutational landscapes has been the analysis of local (typically trinucleotide) mutation signatures. Large-scale pan-cancer analyses revealed an extensive diversity of mutation signatures between and within cancer types^1,24–26^. Some mutational processes are shared (e.g., those manifesting as single base substitution (SBS) signatures 1, 5, and 40), and others are more specific to subsets of cell or cancer types (e.g., MMR deficiency). Previous studies showed that different mutational processes – and their resulting mutational signatures – have differential relationships to replication timing^10,12,15,27,28^. For example, SBS signatures 1, 8, 9, and 17 were shown to be enriched in late replicating regions of the genome, while SBS 5, 21, 40, and 44 showed either bias to early replication or no bias at all. Another property of mutations that we and others have previously described is DNA replicative strand asymmetry, in which certain mutation types tend to occur more often on either the leading or the lagging strands of replication^15,29,30^. Replicative strand asymmetry is characteristic of several mutational signatures (notably SBS 2, 3, 13, and 17), while others are not coupled to asymmetry, e.g., signature SBS 8 is more often observed in late replicating regions but does not show significant replicative strand asymmetry^27^. A further relevant pattern is mutational clustering. For example, clusters of 2-10 mutations caused by the combination of APOBEC3B enzyme activity, replicative errors introduced by DNA Polymerase η, and/or MMR (known as the ‘omikli’ pattern) were shown to be enriched in early replicating regions of the genome, while non-clustered mutation caused by similar mechanisms are late-biased^31,32^.

Previous studies that established how mutational processes relate to DNA replication have assumed that any given process relates to replication timing and strand bias in a constant way. However, it is becoming increasingly clear that mutational processes may be heterogeneous not only in their quantity across cell/cancer types, but also in their relation to replication dynamics across cell types^1,27,28^. This complexity has led to conflicting conclusions among different studies. For example, signature SBS 1 (caused by spontaneous deamination of 5-methylcytosine to thymine) has been reported by different studies to be biased toward early replication, late replication, or neither^10,15,28^. Similarly inconsistent conclusions have been proposed for SBS 5, 40, and others^10,15,28^. These conflicting results could be reconciled if additional, orthogonal factors that vary within and between cell types affect the relationship of mutational processes to DNA replication timing.

Here, we utilized several complementary cell types and hundreds of individuals to perform high-resolution comparisons of mutation rate, pathways, replicative strand asymmetry, and clustering with respect to cell-type-specific replication timing. We first revisit the relationship of mutations and mutational pathways to cell type-specific replication timing patterns. Then, we use two B-cell-types as model systems to identify known and novel factors – and their interactions – that shape the heterogeneity of the mutational landscape with respect to replication timing and strand bias. We discover that global mutation load is broadly associated with the proportion of mutational signatures and their replicative strand asymmetry. We also show that the rate of mutation clustering is associated with the late replication enrichment of a mutational signature. Leveraging these findings, we perform a detailed investigation of mutational pathways on the X-chromosome. Specifically, we perform large-scale mutation phasing to determine if the random and late replication of the inactive X-chromosome influences its mutational landscape. Our results demonstrate that the relationship between the mutational landscape and DNA replication is shaped by a myriad of cell line-specific factors such as mutation load, active mutational processes, mutational clustering, and chromosome inactivation.

## Results

### A catalogue of somatic mutations in five cell types/lines

We called somatic mutations in five cell types/lines for which matched replication timing data is either available or was generated here. These cell types included B-lymphoblastoid cell lines (LCLs), B-cell chronic lymphocytic leukemia (CLL), and three colon cancer cell lines to contrast with the B-cell-related data.

LCLs are Epstein-Barr virus (EBV) -transformed B-cells and are widely available for many individuals. We called LCL mutations by comparing 1662 individuals to their genotyped parents using whole-genome sequence data from six sequencing cohorts (**Table 1, Table S1**). We called 885,655 autosomal single nucleotide variant (SNV) mutations in the offspring by identifying Mendelian errors in parent-offspring allelic inheritance. Autosomal mutation counts ranged from 66 to 8737 per offspring (median 408; 0.169 mutations/Mb) (**Fig 1A, Fig S1A**), consistent with other quantifications of somatic mutations in B-cells^33,34^. We observed two prominent modes and a long tail of mutation count across offspring. This is also consistent with previous mutation calling in the 1000 genomes project (1kGP) offspring and is thought to result from LCL culture age^35^ (**Fig 1A**). Only 0.73% of mutations were functional as predicted by a SNPeff^36^ (4.3t) high or moderate variant impact score. Using monozygotic twins, we estimated the fraction of misidentified parental variants as less than 9.66% (see **Methods**; **Fig S1B-E**). Additionally, we used replicate sequencing of 51 samples to estimate the rate of genotyping errors. We found a median of 93.1% of mutations were supported in samples resequenced once, while 99.8% of mutations were supported at least once in a sample resequenced five separate times (**Fig S1F; Table S1**). Together, mutations in LCL are primarily somatic and reflect LCL biology.

**Table 1.** Mutation data sources. **\*** Further sequencing platform details could not be ascertained.

| Mutation source |  | Number of offspring or samples | Platform | Approx. coverage | Original genome version | Mutation calling method |
| --- | --- | --- | --- | --- | --- | --- |
| LCL | iHART | 1028 | HiSeq X (2 x 150) | 35X | hg19 | Parent-offspring |
|  | 1kGP | 602 | NovaSeq 6000 (2 x 150) | 30X | hg38 | Parent-offspring |
|  | Repeat expansion | 9 | HiSeq X (2 x 150) | 30X | hg19 | Parent-offspring |
|  | Illumina platinum | 13 | HiSeq 2000 (2 x 100) | 50X | hg19 | Parent-offspring |
|  | This study | 12 | HiSeq X (2 x 150) | 15X | hg38 | Parent-offspring |
|  | Polaris | 49 | HiSeq X (2 x 150) | 30X | hg19 | Parent-offspring |
| CLL | CLLE-ES, ICGC | 151 | HiSeq* | NA | hg19 | Tumor-normal |
| HCT116 |  | 6 | HiSeq X (2 x 150) | 15X | hg38 | Passage |
| HT115 |  | 5 | HiSeq X (2 x 150) | 40X | hg38 | Passage |
| LS180 |  | 5 | HiSeq X (2 x 150) | 40X | hg38 | Passage |

**Fig 1.**
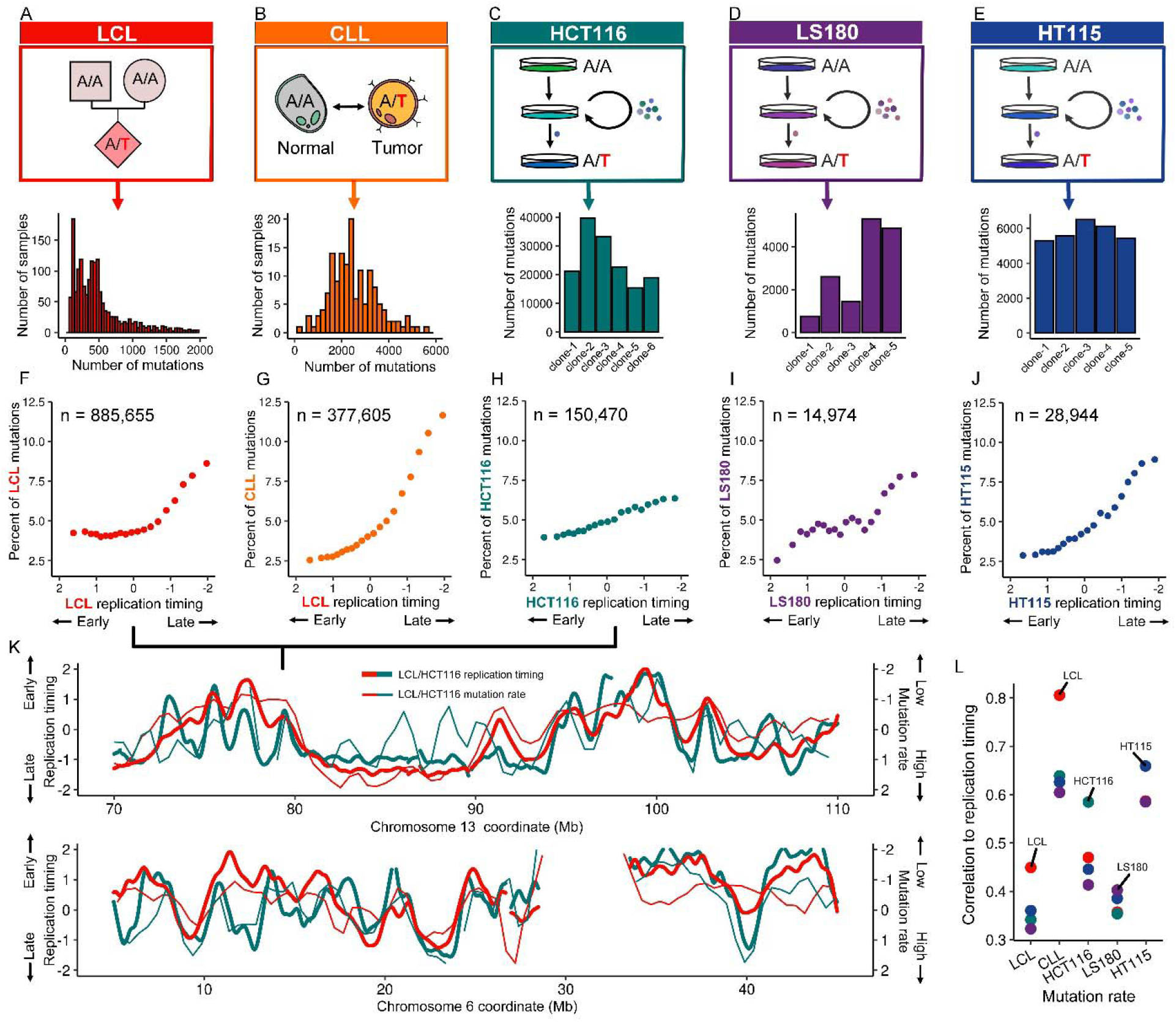
Mutation rate association with DNA replication timing varies in a cell type-specific manner. (A-E) Mutation sources and autosomal counts. (F-J) Autosomal mutation counts in 20 replication timing bins of uniform genome content. (K) Mutation rate correlates to the cell type-specific replication timing in HCT116 and LCLs. Mutation rate is calculated as the mean number of mutations across all samples of the same cell type in a 1Mb sliding window with a 0.5Mb step. Mutation rates are normalized to an autosomal mean of zero and a standard deviation of one to control for the different mutation rates in the two cell types. (L) Mutation rates correlate most strongly with replication timing profiles of the same cells/cell type. Correlation values are Pearson’s correlation coefficients.

To compare LCL mutations to DNA replication timing, we used the same whole-genome sequencing of the offspring to infer replication timing profiles from read depth fluctuations along chromosomes^37,38^. Replication timing is inferred from copy number as early replicating regions have greater read depth in a population of proliferating cells. We then averaged the data for all cell lines to create a single “consensus” LCL replication profiles used for downstream analyses.

To complement the analysis of LCLs, we incorporated mutations derived from 151 CLL patients (**Table 1, Table S1**). CLL is a malignancy of exclusively B-cells, rarely involves EBV infection^39,40^, and has been studied in depth at the genomic level^41^. CLL is a late-onset disease; the mean donor age among samples used in this study was 65.7 years. Tumor-normal mutation calling and filtering identified 377,605 autosomal mutations with a median of 2,368 mutations per patient (0.98 mutations/Mb; range: 221-5629; **Fig 1B**). Of note, due to the primary tumor source of CLL^42^, we could not generate a reference CLL replication timing profile and instead used LCL replication timing to compare to CLL mutations, given that similar cell types have conserved replication timing^43,44^.

As a final point of reference, we incorporated mutational accumulation experiments in three colon adenocarcinoma cell lines. Two cell lines, HCT116 and LS180, possess microsatellite instability (MSI) resulting from loss of functional mismatch repair (MMR). The third, HT115, was microsatellite stable (MSS) with intact MMR. To accumulate mutations, cell lines were sequentially passaged, and single-cell daughter clones were then isolated, expanded, sequenced and compared to the original parental clone (**Fig 1C-E**). Mutations from LS180 and HT115 were sourced from Petljak *et al*., 2019^25^. The cell lines were passaged for 44 and 45 days, respectively, and five daughter subclones were isolated from each line. LS180 yielded 14,974 autosomal mutations (range: 749-5310; median: 2601) and HT115 yielded 28,944 (range: 5296-6511; median: 5,572). HCT116 was passaged by us 100 times (approximately one year) and six daughter subclones were isolated. HCT116 yielded 150,470 autosomal mutations (range: 15,385-39,469; median: 21,846; 9.74 mutations/Mb). Replication timing profiles for LS180 and HT115 were produced by sorting and sequencing G1 and S phase cells^11,21^. An HCT116 mean reference replication timing profile was generated from the whole genome sequencing of the six daughter subclones (this was achievable since HCT116 is near diploid) and further validated by comparison to a profile generated by G1/S sequencing (see **Methods**).

### High resolution comparison of mutation rates to DNA replication timing

Given our large catalog of cell line mutations and the high-resolution analysis they enable, we first sought to refine the relationship of mutation rate to replication timing. We divided the autosomal replication timing profiles into 20 bins of equal genomic proportions organized from the earliest replicating fraction to the latest and counted the number of mutations of each respective cell type within the replication timing range of each bin. While all cell types showed continuous increases in mutation rate with later replication, these relationships differed considerably among cell types (**Fig 1F-J**). Both B-cell-derived cell types, LCL and CLL, showed exponential-like increases in mutation rate from the earliest to latest replicating bins. In LCL, we confirmed the exponential-like relationship independently in the two largest population cohorts (**Fig S1H, I**). Interestingly, LCL only showed an increase in mutation rate in the second half of S-phase, whereas CLL showed a continuous increase (**Fig 1F, G**). CLL demonstrated a more dramatic overall increase in mutation rate, with 4.58-fold more mutations between the latest and earliest replicating bins (from 2.55% of mutations to 11.67%) than LCL (1.90-fold; **Fig 1F, G**). The above differences demonstrate that LCL and CLL mutation landscapes are distinct despite their shared B-cell type. We also observed strong increases in mutation rate in HT115 and LS180, with 3.10-fold and 3.18-fold more mutations in the latest replicating bins than the earliest, respectively (**Fig 1I, J**). In contrast, HCT116 showed a diminished relationship, with an only 1.63-fold (3.90% to 6.35%) increase in mutation rate (**Fig 1H**). The contrast between the cell types, demonstrated most profoundly when comparing CLL and HCT116, establishes a wide disparity in how mutation rates relate to DNA replication timing.

The relationship between replication timing and mutation rates was also apparent visually: plotting mutation rates as continuous profiles along chromosomes revealed a cell-type-specific correspondence with replication timing (**Fig 1K; S1K**). Indeed, the mutation rate in each cell type was most strongly correlated to its matching replication timing profile (**Fig 1L**). Overall, our comprehensive data set comparing mutation rates with matching replication profiles establishes their global correlation but also the heterogeneity among cell types.

### A heterogeneous relationship between replication timing and mutational signatures

To further probe the heterogeneity by which the mutational landscape relates to replication timing, we deciphered the underlying mutational pathways in each cell type and investigated how the rate of each of them varies across the genome in relation to cell type-specific replication timing programs. Specifically, we asked if the disparity in mutation rates between early and late replicating regions could be attributed to specific mutational pathways.

We first determined which mutational processes were active in each cell type and in what proportions. We annotated autosomal mutations in their trinucleotide context and fit COSMIC v3.2 SBS mutational signatures in each cell type. To prevent signature overfitting, we selected a subset of signatures for each cell type based on biologically expected mutational pathways. In CLL, SBS 1, 5, 9 and 40 are established as the predominant mutational signatures^1,28,33,45^. SBS 1, 5, and 40, are clock-like signatures – highly ubiquitous signatures of unknown etiology that increase in abundance with age^1,46^. The proposed etiology of SBS 9 is somatic hypermutation (SHM), a pathway prominent in, and nearly exclusive to, B cells^1,33,34,45^. SHM primarily targets the immunoglobulin heavy chain (*IGHV*) gene but has abundant off-target activity^31,34,47,48^. While, to our knowledge, mutational signature analysis has not been performed in LCL before, we found that the same signatures (SBS 1, 5, 40, and 9) best explained LCL mutations with a cosine similarity of 0.96 for LCLs (compared to 0.97 for CLL). In LCL, it is established that SHM is ongoing after EBV transformation^39,49^. We found that SHM was present globally in both CLL and LCL, but the proportion of mutations explained by SBS 9 was higher in LCL (30.0±0.12% of all autosomal mutations) than in CLL (14.8±0.15%) (**Fig 2A, B; Fig S2A**). Mutations in the MSI cell lines HCT116 and LS180 could be explained by combinations of the six MMR-deficiency (MMRd) signatures: SBS 6, 14, 15, 20, 26, and 44^1^. Along with the common clock-like SBS 1, 5, and 40, we found MMRd signatures SBS 21 and 44 best explained autosomal mutations in both cell lines (cosine similarity of 0.97 in HCT116 and 0.98 in LS180). The MMRd signatures comprised a similar proportion of autosomal mutations in these two cell lines (49.5±0.30% and 47.7±0.95%, respectively) (**Fig 2C, D; Fig S2A**). HT115 is known to have functional mutations in the exonuclease domain of POLE (DNA polymerase ε). The study from which we sourced the HT115 data showed all daughter subclones had additional mutations in the MMR genes *PMS2, MSH6*, and *MSH3*^25^. (One daughter subclone also had a heterozygous POLD1 (DNA polymerase δ subunit) mutation, although it’s signature accounted for a negligible proportion of genomic mutations^25^ and was therefore not further considered in our analysis). SBS 10a-b (POLE mutations), SBS 14 (concurrent MMRd and POLE mutations), SBS 21 (MMRd), and the common clock-like SBS 1, 5, and 40 best explained HT115 autosomal mutations (cosine similarity 0.95). The signatures resulting from POLE mutations and MMRd comprised a total of 53.1±0.63% of autosomal mutations (**Fig 2E; Fig S2A**).

**Fig 2.**
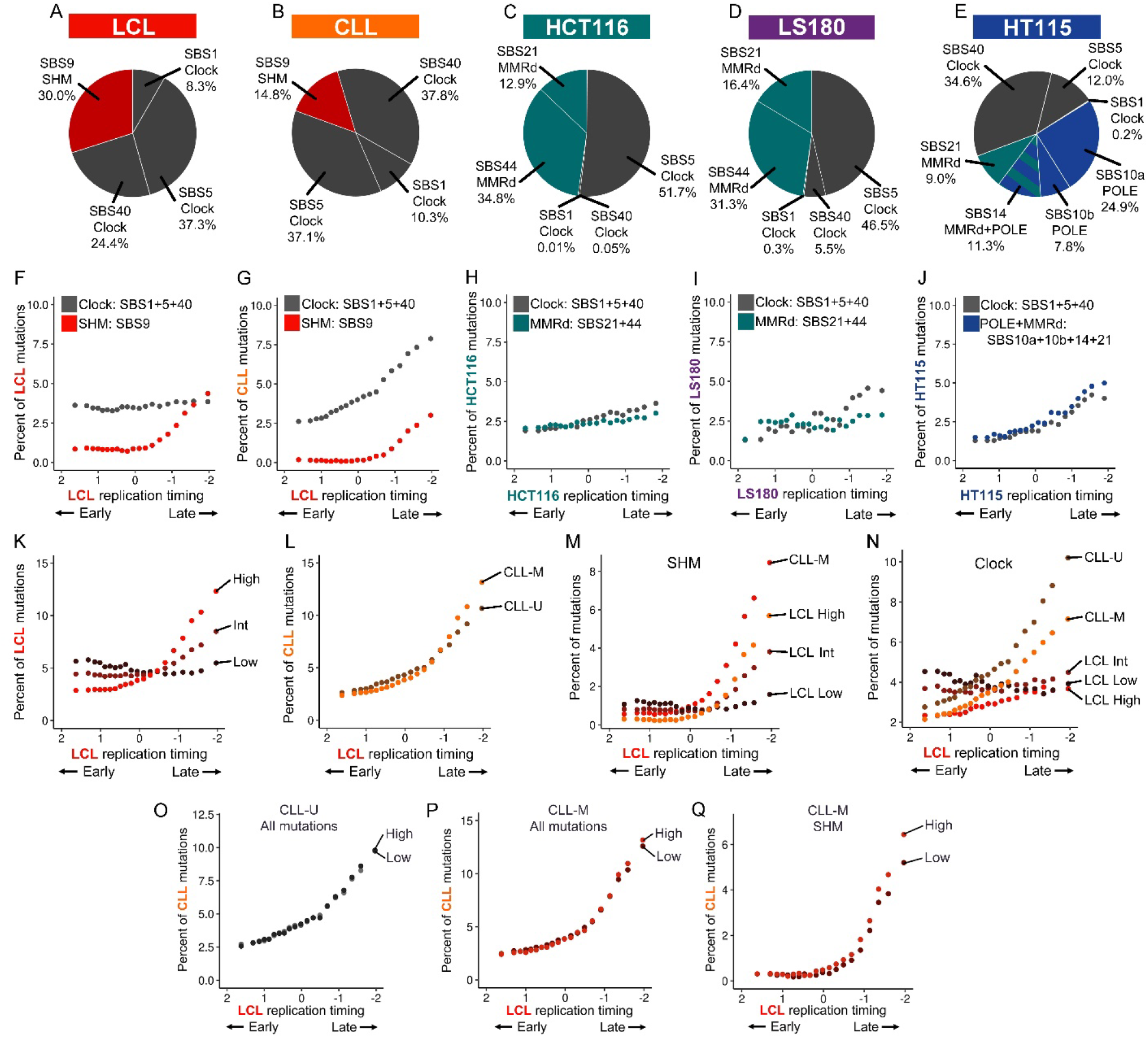
Mutational signatures association with DNA replication timing varies in a cell-type-specific manner. (A-E) Proportion of individual mutational signatures contributing to the total pool of autosomal mutations in each cell type. (F-J) Abundance of mutational signatures in 20 replication timing bins. (K) The relationship of autosomal mutation counts to replication timing in the high, intermediate, and low LCL mutation load groups. (L) The relationship of autosomal mutation count to replication timing in CLL samples stratified by *IGHV* mutation status. (M-N) Abundance of SHM (M) and clock-like mutations (N) as a function of replication timing in the LCL mutation load groups and CLL samples by *IGHV* mutation status. (O) The distribution of total autosomal mutations in CLL-U samples in high and low mutation load groups. (P) As in panel O for CLL-M samples. (Q) The distribution of SHM in the CLL-M high and low groups.

Having established the main mutational signatures contributing to mutations in each cell type/line, we analyzed their relation to replication timing by fitting signatures to mutations in 20 autosomal DNA replication timing bins. We combined the contributions of SBS 1, 5, and 40 into a unified clock-like mutational category, SBS 21 and 44 into an MMRd category for HCT116 and LS180, and SBS 10a, 10b, 14, and 21 in an MMRd+POLE category for HT115.

Several mutational signatures showed distinct relationships to replication timing. In LCL and CLL, SHM (SBS9) contribution increased 16.88- and 5.13-fold, respectively, between the earliest and the latest replication timing fractions (**Fig 2F, G**). In HCT116 and LS180, MMRd contribution increased modestly at 1.60- and 1.09-fold more mutations (**Fig 2H, I**). Compared to SHM and clock-like mutations, MMRd mutations were more uniformly distributed across the genome. This is consistent with previous findings that showed mutations in MSI cancers are less enriched at late replicating parts of the genome^16,50^. In HT115, MMRd+POLE mutations were enriched in late replicating regions in a similar pattern to clock-like mutations, at 2.24x more mutations (**Fig 2J**). Given the stronger replication timing dependence of the combined MMRd+POLE signature compared to MMRd alone, it can be inferred that POLE-derived mutations are specifically enriched in late replicating areas of the genome.

The clock-like category, which explained a substantial proportion of autosomal mutations in all cell types, showed different relationships to replication timing in each cell type. The strongest association was observed in LS180, with 3.42-fold more autosomal mutations in the latest versus earliest replication timing fraction, followed by HT115 (3.12-fold), CLL (3.01-fold), and HCT116 (1.90-fold) (**Fig 2F-J**). In contrast, clock-like mutations showed no apparent relationship to replication timing in LCLs. When considering individual signatures, mutations contributed by SBS 1 – which represents spontaneous deamination of 5-methylcytosine to thymine^1^ – were enriched in late replicating regions in CLL but not in other cell types (**Fig S2B**). SBS 5 and 40 were similarly variable among cell types, although their mutational spectra similarity^1^ precluded associating each of them separately with replication timing. Taken together, the relationship between mutation rates and DNA replication timing varies by mutational pathway and in different ways across cell types.

### Heterogeneity of mutational replicative strand asymmetry

Another property of mutations and mutational signatures that varies along the genome is their tendency to occur on the leading or lagging replicative strands. Extending from the results above, we systematically evaluated the relationships between replicative strand and mutational rates, stratified by mutational signatures and replication timing.

We used the slope of replication timing profiles in each cell type/line to assign replicative strand to mutations (**Fig 3A**): a negative slope on a replication timing profile indicates that the positive genome strand replicates as the leading strand, while a positive slope implies that the positive strand replicates as the lagging strand^30^. Due to uncertainties surrounding the locations of replication origins and termini (peaks and valleys), we regarded 100Kb on either side of a replication direction change as undefined strandedness. While the strand-of-origin of any particular mutation cannot be determined without additional information, the replicative asymmetry of mutations can be evaluated by parsing mutations based on the genomic strand and therefore replicative strand of the substituted pyrimidine base^15,30,51,52^ (**Fig 3A**; see **Methods**). This established approach can determine replicative strand bias based on the ratio of pyrimidine base substitutions. Accordingly, a positive log2-ratio asymmetry value indicates greater leading strand bias of a given mutation type, while negative values indicate greater lagging strand bias.

**Fig 3.**
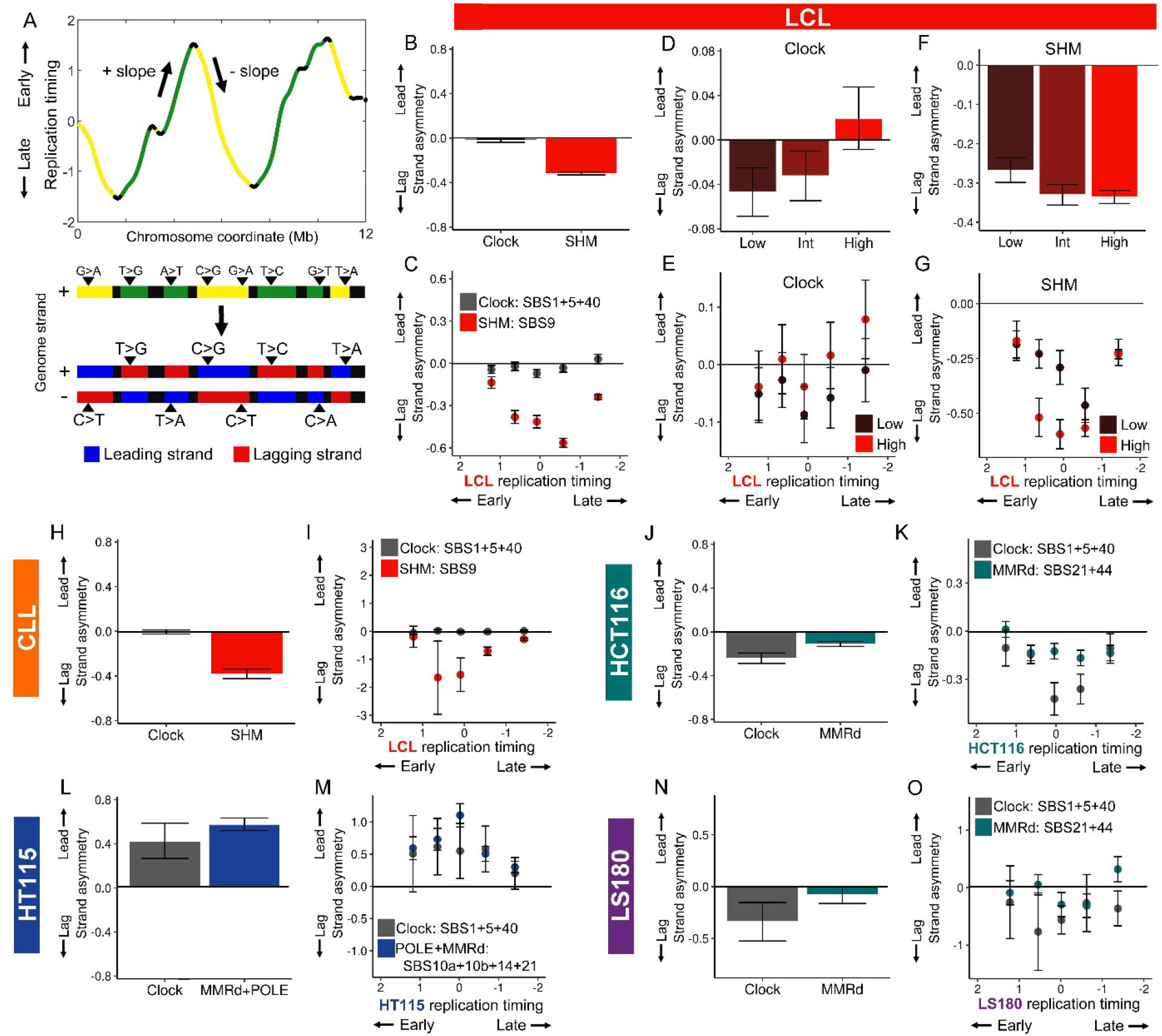
Mutational replicative strand asymmetry varies with replication timing and mutation load. (A) Partitioning mutations by replicative strand. Top: negative slope on a replication timing profile indicates that the positive genome strand replicates as the leading strand, and vice versa for a positive slope. Bottom: Mutations are partitioned to the leading or the lagging strand based on the genome strand and replicative strand of the substituted pyrimidine base. (B) Genome-wide autosomal replicative strand asymmetry for LCL mutational categories. (C) Replicative strand asymmetry for LCL mutational categories in five replication timing bins of uniform genome content. (D-E) Clock-like mutational asymmetry in LCL mutation load groups (D) and as a function of replication timing (E). (F-G) SHM mutational asymmetry in LCL mutation load groups (F) and as a function of replication timing (G). (H-O) As in panels B and C, the replicative strand asymmetry for the mutational pathways in CLL (H-I), HCT116 (J-K), HT115 (L-M), and LS180 (N-O). For all panels, error bars represent the standard error of replicative asymmetry.

We validated strand assignment using four mutational signatures with known replicative strand asymmetries: POLE exonuclease domain mutations result in elevated C>A and C>T mutation on the leading replicative strand^30,52,53^, as indeed we observed for the POLE mutation signatures SBS10a (primarily C>A) and SBS 10b (primarily C>T) being significantly enriched on the leading strand in HT115 (asymmetry values of 0.79±0.07 and 0.73±0.11, respectively; **Fig S3A**); in MMRd, C>T mutations are known to be more abundant on the leading strand^15,54^, consistent with our observation for SBS 44 (MMRd signature characterized by C>T mutations) being enriched on the leading strand (asymmetry value of 0.49±0.03 in HCT116 and 0.57±0.13 in LS180; **Fig S3A**); similarly, T>C substitutions associated with MMRd are more abundant on the lagging strand^30^ and we found SBS 21 (MMRd signature characterized almost exclusively by T>C mutations) to be enriched on the lagging strand (−1.87±0.07 in HCT116, -1.25±0.17 in LS180, and -0.45±0.12 in HT115; **Fig S3A**).

Having demonstrated the effective assignment of replicative strand asymmetry of mutations, we characterized genome-wide replicative strand asymmetry for mutational pathways in the five cell types/lines. Clock-like mutations showed leading strand asymmetry in HT115, yet lagging strand asymmetry in HCT116 and LS180, and no strand asymmetry in LCL and CLL (**Fig 3B, H, J, L, N**). These were surprising results, especially since a previous study that used mutations pooled from many cancer types reported that the clock-like signatures SBS 1 and 5 do not show any strand assymetry^15^. MMRd showed minor lagging strand asymmetry in HCT116 and LS180, which can be explained by the combined abundances and opposing replicative strand asymmetries of SBS 21 and 44 (**Fig 3J, N; Fig S3A**). On the other hand, the POLE+MMRd mutational pathway in HT115 showed substantial leading strand asymmetry, which could be attributed to the overpowering replicative strand asymmetries of POLE mutations over MMRd (**Fig 3L; Fig S3A**). Finally, SHM showed lagging strand asymmetry in LCL and CLL (**Fig 3B, H; Fig S3A**), consistent with previous studies^15,30^.

We next evaluated the replicative asymmetry of mutational pathways with respect to replication timing. Due to the lower number of mutations assigned to a given strand, we analyzed five instead of 20 genomic bins. Replicative strand asymmetry of clock-like mutations did not change between the replication timing fractions in all cell types except for HCT116, where greater lagging strand asymmetry was evident in the middle replicating fractions (**Fig 3C, I, K, M, O**). Thus, as with mutations in general (above), the relationship of the clock-like category to replication timing was variable across cell types/lines. Lagging strand asymmetry for MMRd mutations in HCT116 and LS180 also did not change between replication fractions (**Fig 3K, O**). However, the asymmetry for the individual MMRd signatures SBS 21 and 44 showed the strongest lagging and leading strand asymmetry values respectively in the middle replicating fractions (**Fig S3B**). A similar trend was observed for SHM and POLEd+MMRd (**Fig 3C, I, M**). This mid-S-phase pattern of greater asymmetry was found in the individual signatures SBS10a, 10b, and 14 (**Fig S3B**). By removing 500Kb regions flanking slope directionality changes, we ruled out that these mid-S enrichment patterns were due to uncertainty in calling replication origin and terminus locations and hence replication direction in their vicinity (**Fig S3C**). Taken together, mutational signatures and pathways showed variable replicative strand asymmetry patterns with respect to replication timing. Importantly, these cell-type-specific asymmetry patterns were distinct from the mutation rate patterns described above. More generally, our analyses so far reaffirm and extend previous findings that the relationship between mutational pathways and replication timing is heterogeneous across cell types and provide a foundation for the more detailed investigations to follow.

### Mutation load and SHM modulate the mutational landscape

Having demonstrated variability in how mutation rate fluctuations relate to replication timing, we sought to identify additional factors that differ between and within cell types and that could further account for such heterogeneity. For this, we focused on LCL and CLL due to their inclusion of multiple samples and shared mutational pathways. A major difference between these two cell types is the elevated mutation load (also known as mutation burden) of CLL, as defined by the total number of autosomal mutations per sample (**Fig 1A, B**). We thus asked if mutation load itself relates to the distribution of mutations with respect to replication timing. To test this, we began by dividing the LCL offspring (which were more numerous than the CLL samples available here; we return to CLL below) into three groups based on the number of autosomal mutations, such that each group contained a similar (∼295,500) total number of mutations (**Fig S4A**). A “low mutation load” group contained ≤489 mutations per offspring (1066 offspring); a “high mutation load” group had ≥1104 mutations per offspring (174 offspring); and an “intermediate mutation load” group contained the remaining 422 offspring. We observed that the relationship of mutation rate to replication timing was substantially more pronounced in the high mutation load group, with 4.17-fold more mutations in the latest replicating fraction than the earliest (**Fig 2K**). In comparison, the intermediate mutation load group showed a less dramatic increase with 1.85-fold more mutations in the latest fraction, while the low mutation load group did not show enrichment at all for mutations in late replicating parts of the genome (0.98-fold difference). Importantly, this result was not attributed to statistical power, as all groups had a similar and sufficient number of mutations analyzed. This pattern was also evident for individual offspring, where greater mutation load corresponded to consistently later replication timing bias, including when offspring were down sampled to only 80 mutations to control for possible power differences among samples (**Fig 4A-C**). Thus, LCLs with a greater number of autosomal mutations exhibited an inherently stronger enrichment of mutations in late-replicating genomic regions.

**Fig 4.**
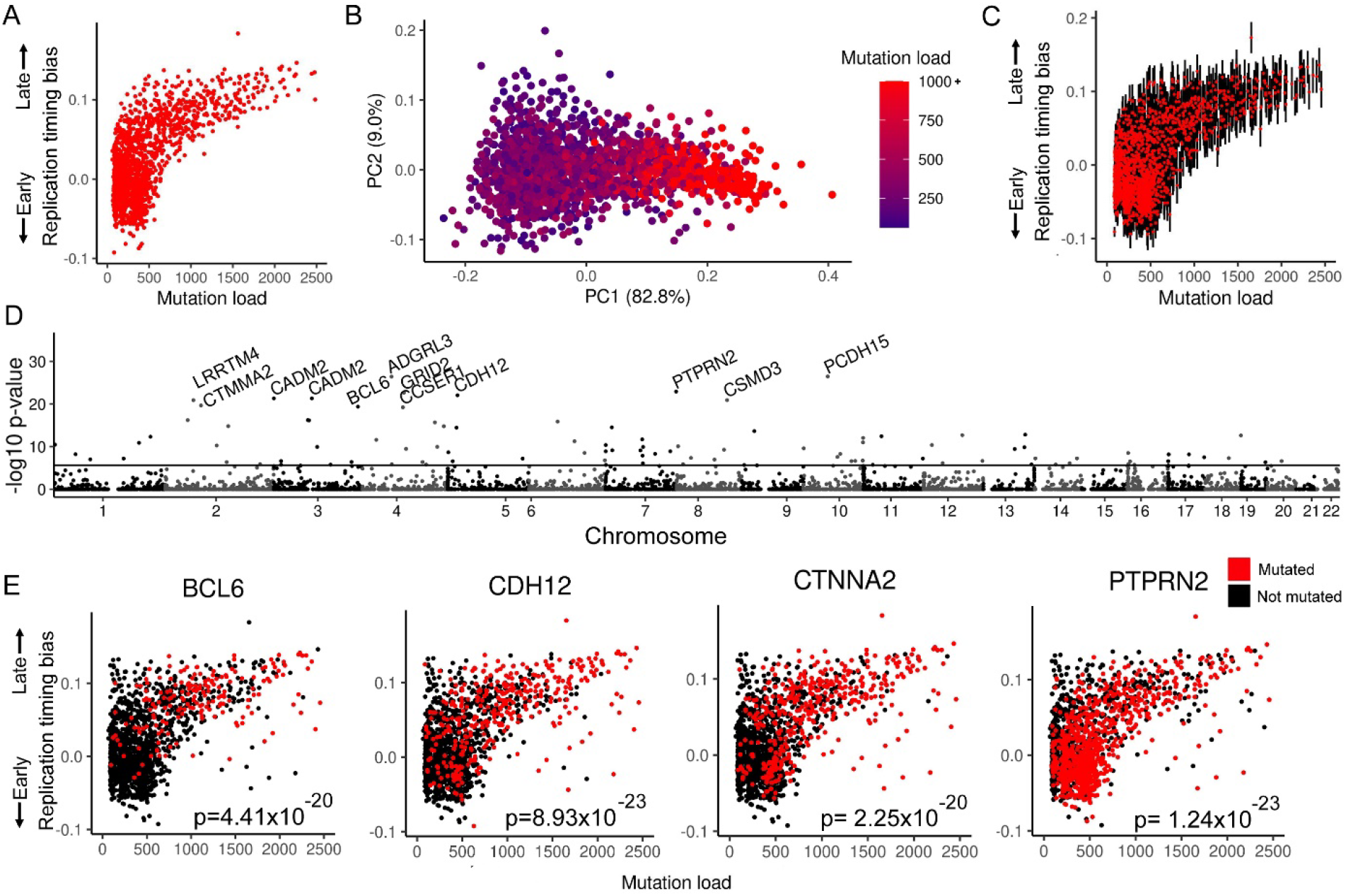
Individual LCL late replication timing bias and candidate gene associations. (A) Replication timing bias, calculated as the linear slope of mutation percentages in four replication timing bins, increases with mutation load across individuals. (B) PCA of the percentage of mutations in four replication timing bins calculated for panel A. PC1 corresponds to mutation load. (C) Down sampling of individual LCL samples to 80 genome-wide mutations. Red dots indicate the mean slope of 1000 iterations of samplings for each mutation load. Error bars represent the standard deviation of samplings. (D) Association of mutated gene frequency to late replication timing bias of individual samples (as shown in panel A) corrected for mutation load. Black line indicates the Bonferroni-corrected p<0.05 divided by number of tested genes. The top 11 most significant genes are highlighted. (E) Selected genes from panel D showing mutation status in individual LCLs.

We asked if these differences between mutation load groups are related to particular mutational signatures. Accordingly, we fit SHM and clock-like mutational signatures to the stratified LCL mutation load groups. We found that the proportion of mutations attributed to SHM decreased from 43.46±0.22% of mutations in the high mutation load group to 25.74±0.19% and further down to 21.01±0.18% in the intermediate and low mutation load groups, respectively. This trend was also observable in individual samples, as SHM contribution correlated, albeit modestly, with mutation load (Pearson’s *r* = 0.34, p<1×10^−16^). Therefore, the high global mutation count in LCLs is disproportionately driven by SHM. With respect to replication timing, the high mutation load group showed the greatest enrichment in late-replicating regions for both SHM and the clock-like category, with 15.1-fold and 1.57-fold more mutations in the latest replicating fraction compared to the earliest, respectively (**Fig 2M, N; Fig S4B**). This relationship was less pronounced in the intermediate mutation load group, with a 4.69-fold increase in SHM abundance and a 1.22-fold increase in clock-like abundance. The low mutation load group showed enrichment for neither SHM nor clock-like mutations in late replicating regions of the genome (**Fig 2N**). Together, these findings indicate that the distribution of mutations, most prominently of SHM origin, varies in LCLs in accordance with mutation load.

CLL samples provided an opportunity to further investigate how mutation load and signature proportions shape the mutational landscape. Since CLL comprises two subtypes that differ by the mutational status of *IGHV* and therefore by mutation load, we first separated CLL samples by subtype. CLL tumor samples with a mutated *IGHV* (CLL-M) are known to have undergone SHM, and patients have a higher survival rate than those with an unmutated *IGHV* (CLL-U)^55^. The CLL samples used in this study included both CLL-M and CLL-U, but the *IGHV* mutation status of individuals was unreported. We therefore devised a way to use mutational signature analysis as an alternative means of inferring SHM activity and thus CLL subtype. Accordingly, we fit the CLL mutational signatures (SBS 1, 5, 9, and 40) to the autosomal mutations in individual samples. We assigned 80 samples with a consistent >2% SHM contribution over 1000 bootstrap samples as CLL-M, and another 68 samples with a consistent 0% SHM contribution as CLL-U (**Fig S4C**). Three remaining samples were ambiguous and not analyzed further. The CLL-M group contained a median of 2,620 autosomal mutations per sample (216,451 total mutations; **Fig S4D**), while the CLL-U group contained a median of 1,986 autosomal mutations per sample (138,113 total mutations). This was a significant difference in mutation burden between the two CLL subtypes (two-tailed t-test: p = 1.63×10^−5^). In CLL-M samples, a median of 25.4±0.04% of all mutations (591 mutations per sample) were contributed by SHM, which can fully account for their increased global mutation count.

Mutations in CLL-M and CLL-U samples showed exponential-like increases with replication timing (**Fig 2L**). This effect was slightly stronger in CLL-M (5.54-fold more mutations in the latest replicating fraction than the earliest) than in CLL-U (4.05-fold). More specifically, in CLL-M, as in LCLs, SHM contribution was greatly enriched in late replicating regions, with 18.9-fold more mutations in the latest replicating fraction than the earliest (**Fig 2M; Fig S4F**). This distribution of SHM mutations in CLL-M comprised the strongest enrichment of mutations in late replicating regions that we observed in all our analyses so far. For clock-like mutations, CLL-M and CLL-U showed similar replication timing relationships with 3.32- and 3.69-fold more mutations, respectively, in the latest replicating fraction than the earliest (**Fig 2N**).

Having CLL subdivided by *IGHV* mutation status, we could then compare high and low mutation load (as for LCL above). We divided CLL-M and CLL-U into two groups each, based on autosomal mutation load. CLL-M samples with higher mutation loads (28 samples with ≥3,011 mutations) showed greater enrichment for all mutations in late replicating regions (**Fig 2P**). Among CLL-M samples, higher mutation load corresponded to greater SHM contribution (20.6±0.30% versus 25.24±0.32%) and greater SHM enrichment in later replicating regions (**Fig 2Q**). CLL-U did not show a pronounced change in mutation enrichment in late replicating regions based on mutation load (**Fig 2O**), likely due to the diminished variability in mutation load among CLL-U samples (**Fig S4D**). Thus, we again observe that the distribution of SHM mutations varies in accordance with mutation load.

We next asked if the influence of global mutation load on the mutational landscape extends to replicative strand asymmetry. We used the stratification of LCL offspring by autosomal mutational load and reevaluated strand asymmetry for the clock-like and SHM mutational categories. There was substantial lagging strand asymmetry for the low mutation load group for clock-like mutations, and a more modest leading strand asymmetry for the high mutation load group (**Fig 3D**). SHM mutations also showed pronounced differences, but with greater genome-wide lagging strand asymmetry in the high mutation load group compared to the low mutation load group (**Fig 3F**). With respect to replication timing, while there were no significant differences between groups for clock-like mutations (**Fig 3E; S3D**), SHM asymmetry differed considerably across the mutation load groups although only within the middle fractions of replication timing (**Fig 3G; Fig S3D**). Specifically, in the middle replicating quintile, lagging strand asymmetry was greater in the high mutation load group. Thus, while SHM contribution to LCL mutations was more pronounced in late replicating regions, lagging strand asymmetry appeared to increase more in mid-S replicating regions with higher mutation load.

Taken together, we identified global mutation load as a novel cell line-specific factor that associates with the distribution of mutations along the genome and with respect to replication timing. In both LCL and CLL-M, elevated mutation load corresponded to increased SHM abundance genome-wide and in late replicating regions specifically. This finding has important implications for interpreting how mutation signatures relate to DNA replication timing, as these relationships may vary based on the mutation loads of individual samples.

A natural explanation for the association between mutation load and replication timing bias is that mutation of a *trans*-acting factor elevates late replication timing bias, and this factor is more frequently mutated in high mutation load samples (either as a direct cause of their high mutation load, or in association with the elevated number of mutations). We tested this in LCLs by associating mutations at the level of genes with individuals’ mutational late replication timing bias, while controlling for mutation load (**Fig 4A**). It is essential to control for mutation load as the nominal number of mutations in any region would be higher with greater mutation load irrespective of replication timing dynamics. We identified several candidates significantly associated with late replication mutational bias, including several linked to cancer risk such as *CSMD3* and *CTNNA2* (**Fig 4D,E**). Of particular interest was *BCL6* (B-cell lymphoma 6), a transcription factor that promotes proliferation of B-cells after the onset of SHM by repressing genes that would otherwise arrest the cell cycle as a result of elevated DNA damage^56^.

We identified 345 mutations within the *BCL6* gene among 192 of the 1662 LCLs. In the high mutation load group, *BCL6* mutations were found in 52.3% of samples compared to only 17.8% and 2.1% in the low and intermediate mutation load group, respectively. This could not be explained by differences in sample mutation load, as high mutation load samples had on average 6.1-fold more mutations than low mutation load samples whereas *BCL6* mutations were 24.9-fold more common. We additionally found *BCL6* mutations in 20.7% of the 906 samples with a late replication timing bias (**Fig 4A**) compared to 5.7% among samples with early or no replication timing bias. Differences in sample mutation load was again ruled out, as samples with a late replication timing bias had on average 1.58-fold more mutations globally whereas *BCL6* mutations were 3.63-fold more common. Mutations in the *BCL6* gene were also found in 26.5% of CLL samples and were far more common in CLL-M (48.8% of samples) than CLL-U (1.5%). Of note, *BCL6* is a COSMIC (v96) census driver of CLL^57^ though our results suggest this gene is more important for CLL-M.

Functional mutations of *BCL6* were rare (as with all genes) as only two were discovered in LCL and one in CLL, though other mutations may still affect the regulation of *BCL6*. An attractive possibility is that *BCL6* mutations arise in LCL culture and promote both a higher mutation load as well as an altered mutational landscape manifesting in late replication mutational bias. Moreover, such mutations may be selected for during LCL culture, consistent with their higher prevalence in older cell lines (although we cannot discriminate between mutation load and culture age as being causally linked to *BCL6* mutation prevalence). If this were the case, *BCL6* could be the equivalent of *BCOR* (*BCL6* corepressor) mutations that are selected for in iPS cell culture^58^; indeed, *BCOR* functions together with BCL6 to repress cell cycle arrest in cells with active SHM. Further research will be required to characterize the role of *BCL6* (and other genes) in the proliferation and mutational landscape of LCLs.

### SHM entails two mutational modes with distinct replication timing and clustering

SHM initiates with the deamination of cytosine into deoxyuracil via activation-induced cytidine deaminase (AID) operating on ssDNA^59,60^. Left unrepaired, C>U deamination converts to C>T mutations during DNA replication^61^. Alternately, the initial deamination can be repaired by non-canonical MMR, which includes DNA synthesis by the low fidelity DNA polymerase η (POLH)^31,61^. POLH synthesis produces proximal A>G and A>C substitutions, the characteristics of SBS 9 and therefore SHM^1,62^. It has previously been shown that a subset of SHM-context mutations in B-lymphocyte cancers (T>C and T>G substitutions with a 3’ A or 3’ T context) cluster at promoters and enhancers of actively transcribed genes and are enriched within 100bp of C>N mutations^31^. Additionally, pooling mutations of SHM origin across many cancer types showed that non-clustered mutations are more enriched than clustered mutations in late replicating regions^31,32^. This indicates that a given mutation pathway, like SHM, could entail distinct mutational modes, each with different relationships to replication timing and other genomic features. It is also conceivable that the presence of such modes would differ across cell types, potentially explaining why SHM is more enriched in late replicating regions in CLL than in LCL.

To test the role of SHM clustering in determining late replication bias, we clustered SHM-context mutations in LCL and CLL by considering two or more SHM-context mutations falling within 500bp of each other as a cluster. We identified 26,759 such clusters in LCLs and 2,624 in CLL, encompassing 37.01% and 7.50% of total SHM-context mutations, respectively. Although there was a nominal increase in cluster number and proportion with replication timing (**Fig S5A-D)**, when controlling for the correlation of mutation rates with replication timing (see **Methods**) we found that, first, clustering in LCL and CLL was significantly elevated (p<1×10^−100^) in every replication timing fraction (**Fig S5A-D**), and second, clustering was relatively more abundant in early replication timing fractions (**Fig S5E-H**). Reciprocally, non-clustered mutations were more abundant in late replication timing fractions (**Fig S5I, J**). Reduced SHM mutation clustering in CLL thus relates to their greater bias towards late replication.

When controlling for gene content across replication timing fractions (and considering each mutation within clusters individually), we found that clustered mutations were significantly closer to genes compared to non-clustered mutations, in both LCL (p<1×10^−246^) and CLL (p<1×10^−55^). This was reminiscent of the gene-enriched *omikli* pattern of cancer mutation clusters^32^. Because genes and clustered mutations are both enriched in early replicating regions of the genome, we compared gene proximity in replication timing bins, controlling for gene content. For the earliest replicating 75% of the genome, clustered mutations in LCL and CLL were significantly more proximal to genes (p<1×10^−10^) than non-clustered mutations (**Fig S5K-N**). Surprisingly, in the latest 25% of the genome, we observed the opposite pattern with non-clustered mutations significantly more proximal to genes (p<1×10^−10^). A yet distinct pattern was observed with regards to C>N mutations, which in the latest replicating fractions were closer to clustered mutations than they were to non-clustered mutations (**Fig S5O-R**). The differing distributions of clustered and non-clustered mutations in relation to genes and C>N mutations further support the notion that there are two distinct SHM mutational modes, representing more than one mutational mechanism that would otherwise be grouped together.

### Unique mutational processes on the inactive X-chromosome

We described above multiple factors that shape, in a cell-type-specific manner, how mutations accumulate along the genome and with respect to replication timing: replication timing patterns; different mutational processes (as manifested in mutational signatures) and their replicative strand asymmetries; and mutation clustering. Individual, cell line-specific factors such as global mutation load further influence the mutational landscape including the extents of late replication bias and replicative strand asymmetry. As a case in point, we examined these factors from the perspective of the unique biology of chromosome inactivation. The inactive X-chromosome (Xi) in females replicates late in S-phase with no discernable replication timing pattern^63^, which is distinct from the active X-chromosome (Xa), the male X-chromosome, and autosomes. This, and the tight link between replication dynamics and the mutational landscape led us to predict that the Xi would also have unusual mutational properties. Consistently, in some cancers, Xi has been inferred to have a higher mutation rate than Xa and the male X-chromosome^8,64^. In our female LCL offspring and CLL samples, we also found that the X-chromosome demonstrated significantly higher mutation rate than autosomes (**Fig S6A, B**). Interestingly, the female X-chromosome also showed a significantly greater abundance of SHM compared to autosomes (**Fig S6C, D**; see further below).

The large-scale, family-based configuration of our LCL samples provides unprecedented power to phase mutations and separately investigate the mutational landscapes of Xa and Xi. This is in contrast to previous studies that investigated Xi mutations by male-female comparisons or with limited expression-phased mutations^8,64^. Xi has been to shown to be clonally propagated^65–67^ and is therefore expected to be detectable in at least a subset of the 746 female LCL offspring. While phasing inherited variants enables discriminating parental chromosome pairs, functional data is required in order to identify the inactive X-chromosome. To this end, we devised an approach using the replication timing data itself, as inferred from sequencing read depth: due to its later replication, the Xi is expected to demonstrate a significantly lower median copy number compared to the Xa (**Fig 5A**). Indeed, female X-chromosomes showed greater parental copy number disparity than autosomes, which we used as a benchmark for assigning X-chromosome identity (specifically, for samples with greater than the 95^th^ percentile disparity on chromosome 14 – the autosome with the closest number of phaseable inherited variants to the X-chromosome; **Fig 5B, C**). This approach yielded reproducible Xi assignments in 17 of 17 replicate sequenced offspring for which assignments could be made. In addition, paternal Xi identity for NA12878 was consistent with RNA expression analyses^68,69^ and with our previous classification for this cell line^63^. Thus, the inactive X-chromosome can be identified, and mutations it harbors can be called, from the same genome sequence data. Accordingly, we identified the Xi in 542 of 746 female offspring (72.65%), of which 293 were paternally X-inactivated and 249 were maternally X-inactivated (**Fig 5D**).

**Fig 5.**
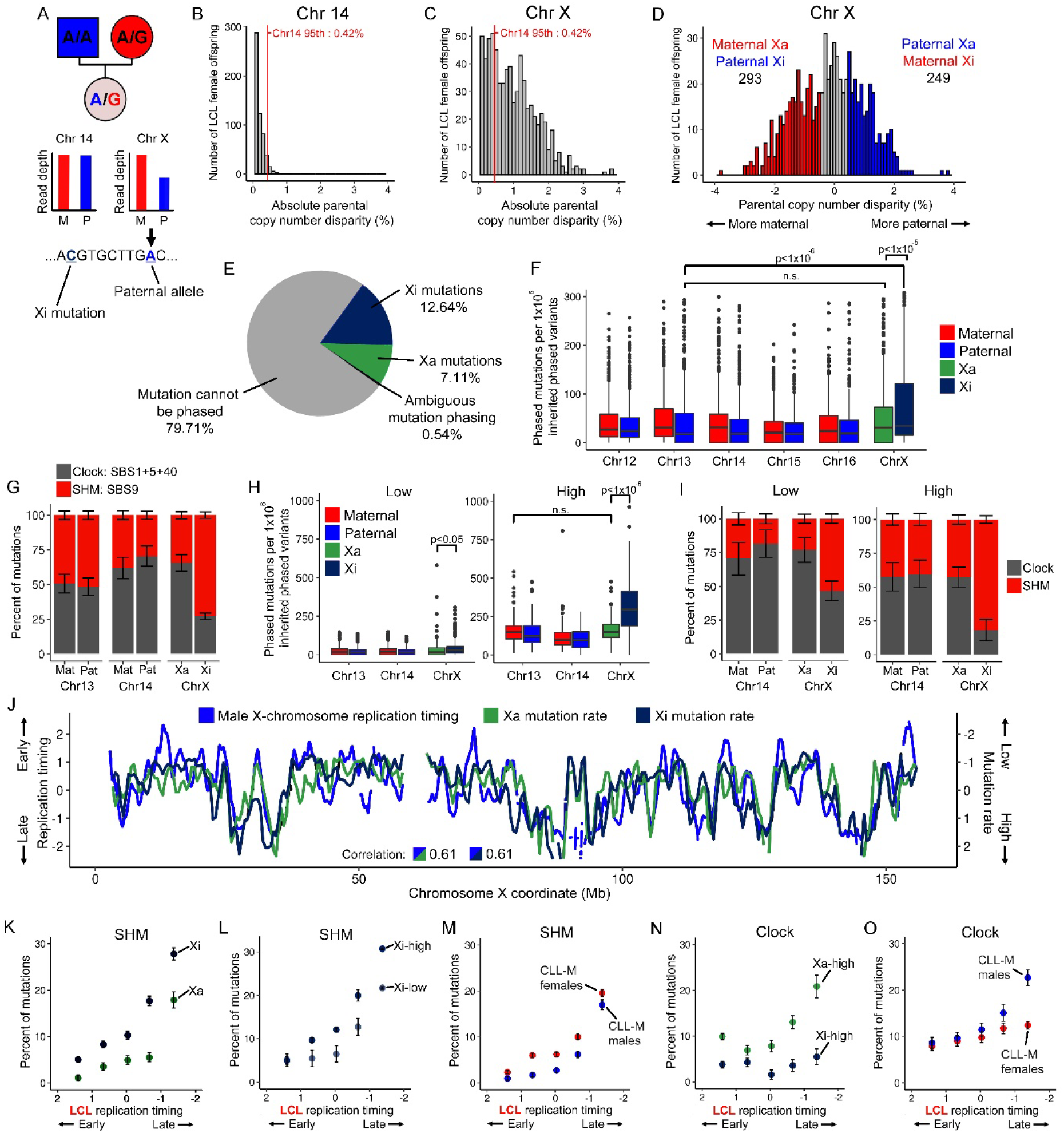
Unique mutational processes on the inactive X-chromosome. (A) Identification of Xi parental identity and mutation phasing. (B) The absolute parental read depth disparity in LCL female offspring on chromosome 14. Disparity was calculated as the absolute difference of paternal and maternal median read depth of inherited phaseable variants divided by their combined median depth. (C) The elevated absolute parental read depth disparity on the X-chromosome in female LCL offspring. Xi was identified in females with a disparity greater than the 95^th^ percentile value from chromosome 14. (D) Xi parental identity classification among females with an identifiable Xi as described in panel (C). Xi is the parental homolog with the lower read depth. (E) The number of phased X-chromosome mutations in females with an identifiable Xi. (F) Xa and Xi mutation rate compared to maternal and paternal homologous autosomes with the most similar number of inherited phaseable variants to chromosome X. Mutation rate was calculated as the number of phased mutations normalized by the number of inherited phaseable variants on each chromosome homolog pair. P-values were calculated from a two-tailed t-test. (G) Proportions of mutational pathways on maternal and paternal homologous autosomes and Xa/Xi. (H) As in panel (F), the mutation rate of phased mutations in high and low autosomal mutation load groups. (I) As in panel (G), the proportions of mutational pathways in high and low autosomal mutation load groups. (J) Pearson correlations of Xa and Xi regional mutation rate (calculated as in Fig 1K and further normalized by the number of inherited phaseable sites in each window) to male X-chromosome replication timing. (K-O) Abundance of mutational pathways on the X-chromosome in five replication timing bins: SHM abundance for Xa/Xi mutations (K), Xi mutations in the high and low autosomal mutation load groups (L), and CLL-M male and female patients (M); Clock-like mutation abundance for Xa/Xi mutations in the high autosomal mutation load groups (N) and CLL-M male and female patients (O). In all panels, error bars represent the standard error of signature fit.

Being able to phase the X-chromosomes across a large set of cell lines, we systematically quantified how mutation rate and mutational processes differed between Xa and Xi. We phased mutations by identifying mutant alleles on the same sequencing read or mate-pair as a phaseable inherited variant (**Fig 5A**). Among the 542 females with an identifiable Xi, we phased 6005 (19.75%) X-chromosome mutations, of which 3844 (64.01%) were assigned to the Xi (**Fig 5E**). This comprises, to our knowledge, the largest collection of Xi- and Xa-parsed mutations. We confirmed that the mutation rate of Xi was 1.78-fold higher (p<1×10^−5^) than that of Xa and significantly higher than any autosome (p<1×10^−6^) (**Fig 5F; Fig S6E**); the mutation rate of Xa was not significantly different from that of autosomes (**Fig 5F**). With regards to mutational processes, the proportions of mutations explained by SHM (34.36±2.49%) and the clock-like mutational category (65.64±5.94%) were similar between the Xa and autosomes (**Fig 5G; Fig S6F**). On the Xi, however, only 27.16±2.38% of mutations were attributable to the clock-like category, while 72.84±2.27% were attributable to SHM (**Fig 5G**). The elevated mutation rate on the Xi can thus be predominantly attributed to SHM.

Given our observation that mutation load relates to SHM enrichment in late-replicating genomic regions, we hypothesized that increased overall mutation load in a cell line would correspond to disproportionately greater Xi mutation rate and SHM abundance. We split the 542 LCL offspring with an identifiable Xi into a low mutation load group with less than 832 autosomal mutations (433 offspring), and a high mutation load group (remaining 109 offspring). Each group contained approximately 157,000 autosomal mutations. As predicted, X-chromosome mutations were proportionally more abundant in the high mutation load group, comprising 11.10% of mutations compared to 8.25% in the low mutation load group. Using phased mutations, we further found that 67.33% of X-chromosome mutations in the high mutation load group were located on the Xi, compared to only 58.14% in the low group (**Fig 5H**). As a control, Xa showed the same mutation rate as autosomes in both groups (**Fig 5H**). This confirms that Xi have an elevated mutation load compared to Xa or autosomes. As further hypothesized, we found that SHM abundance on the Xi was strongly elevated in the high mutation load group, at 81.72±2.71% of Xi mutations compared to 53.37±3.44% in the low mutation rate group (**Fig 5I**). In addition, SHM abundance on the Xi was higher than on the Xa, comprising 38.92% more mutations on Xi than Xa in the high load group, compared to 30.33% in the low group. Taken together, X-chromosome inactivation is associated with an elevated mutation load driven by SHM, thus creating a distinct mutational landscape on the Xi; This disparity of mutation load and SHM composition relative to the Xa is particularly pronounced in cell lines with a greater global mutational load.

### Association of mutational pathways with X-chromosome-specific replication programs

We showed above that the elevated mutation load and SHM abundance on Xi were consistent with its late replication. We next investigated how mutations relate to the random replication pattern of the Xi. If replication timing is a direct modulator of mutation rate, the random replication of Xi would predict a random, uniform distribution of mutations. Using the 542 LCL offspring with an identifiable Xi, we assessed regional mutation rates of phased mutations in 1Mb sliding windows with a 0.5Mb step. As expected, for the Xa, regional mutation rate correlated to male X-chromosome replication timing (*r=*0.61) at similar levels as phased autosomal mutations to autosomal replication timing (**Fig S6G**). Unexpectedly, regional Xi mutation rate demonstrated an equally high correlation to male X-chromosome replication timing (*r=*0.61; **Fig 5J; Fig S6G**). This suggests that Xi mutation distribution follows the ordered replication timing pattern of Xa rather than the random pattern of Xi.

Given the unanticipated result of ordered Xi mutations in LCL, we sought to validate these findings in CLL. Although we were unable to similarly phase CLL mutations, we compared X-chromosome mutations across male and female patients to estimate the mutational landscape of Xi. For autosomes, regional mutation rates in males and females near-equally correlated to replication timing (**Fig S6H**). However, in contrast to LCLs, this correlation was reduced for X-chromosome mutations in female CLL patients (*r=*0.67 among females, 0.76 among males; **Fig S6H**). A principal difference between LCL and CLL is *IGHV* mutation status. As described above, CLL-U mutations are only contributed by the clock-like category, while CLL-M and LCL mutations are partly contributed by SHM. By analyzing CLL-M and CLL-U separately, we found that the correlation for X-chromosome regional mutation rate in CLL-U female patients (*r=*0.46) was distinctively diminished compared to CLL-U males (*r=*0.70) and autosomes (**Fig S6I**). This level of reduced correlation was not observed in CLL-M females (**Fig S6J**). As CLL-U samples lack SHM, we suspected that clock-like mutations are randomly distributed on the Xi while SHM mutations follow more closely the Xa replication pattern.

To study the distribution of SHM mutations on the Xi, we split phased mutations into five bins based on the male X-chromosome replication timing. If SHM mutations are randomly distributed on Xi, we would expect the phased Xi mutations to be distributed independently of replication timing. However, in LCLs, Xa and Xi mutations showed similarly high enrichment for SHM in late replicating regions of the male X-chromosome (**Fig 5K**). Late replicating timing enrichment was stronger for Xi mutations in the high (6.21-fold more) versus low (4.28-fold) autosomal mutation load groups (**Fig 5L**). Thus, the disordered replication timing of Xi does not directly relate to SHM mutation rate in LCLs. To validate this in CLL-M, we expected to observe equal enrichments for SHM in late replicating regions in male and female patients. We indeed found that female CLL-M X-chromosome mutations were similarly enriched in late replicating regions (10.41-fold) as males (12.29-fold; **Fig 5M**). Thus, in both LCL and CLL, Xi SHM mutations distribution follows the ordered pattern of Xa replication timing.

Last, we examined clock-like mutations on the Xi, focusing specifically on the LCL offspring with high autosomal mutation loads (since we only observed late-replication enrichment of clock-like mutations in those; see **Fig 2N**). We found that Xa clock-like mutations in the high load group were enriched in late replicating regions of the male X-chromosome (2.11-fold; **Fig 5N**). However, in contrast to SHM, Xi clock-like mutations were more uniformly distributed with respect to male X-chromosome replication timing (0.99-fold; **Fig 5N**). This supported the hypothesis that clock-like mutations are randomly distributed on Xi. We again validated these results in CLL-M: if Xi clock-like mutations are randomly distributed, we would expect a more uniform distribution of clock-like mutations with respect to replication timing in female versus male CLL-M patients. As anticipated, CLL-M females demonstrated a striking reduction of clock-like mutations in late replicating regions of the male X-chromosome (1.57-fold) compared to CLL-M males (2.63-fold; **Fig 5O**). Taken together, both LCL and CLL suggest that the replication pattern of Xi may directly relate to clock-like, but not necessarily SHM, mutations.

## Discussion

In this work, we sought to identify factors that explain how mutation rate fluctuates with replication timing and how this relation varies across samples. We first affirmed that the relationship between mutation rates and replication timing was heterogeneous by comparing five cell types. We further characterized this variability through the specific mutation signatures of the cell type and found both signature quantity and its replicative strand asymmetry vary in relationship to replication timing. For example, SBS9 was highly enriched in late replicating regions of the genome whereas its asymmetry was most apparent in mid S-phase. Clock-like mutations were distributed more flatly on the chromosome with less prominent asymmetry though these properties varied considerably by cell type. We next showed that individual mutation load and mutation clustering greatly influence the late replication timing bias of mutations, particularly of SHM origin. Greater mutation load corresponded to elevated SHM late replication bias whereas clustered mutations were relatively enriched in early replicating regions. We then uncovered a unique mutational landscape of the inactive X-chromosome, showing Xi contained a higher mutation load explained by elevated SHM activity. We additionally found elevated autosomal mutation load exacerbates the disparity of mutation load and SHM abundance between Xa and Xi. Finally, by comparing the landscape of mutational signatures on Xi, we found evidence for clock-like mutations being directly modulated by replication timing, while SHM mutations are seemingly not. Together, the presence of multiple factors influencing the mutational landscape challenges our understanding of how mutational pathways relate to replication timing.

An unexpected finding was that an individual sample’s mutation load greatly influences whether mutational signatures are enriched in late replicating regions and/or show replicative strand asymmetry. We confirmed this observation among individual LCLs, through the down sampling of LCL mutations, and in CLL, where mutations were identified using a different methodology. The effect of mutation load may largely underly the conflicted reporting of mutation signature quantity and replication timing enrichment across cell/cancer types. For example, a collection of high mutation load LCLs would produce different conclusions about SHM or clock-like category abundance than a collection of low mutation load LCLs. More generally, a lower mutation load cohort may suggest the distribution of a signature is flatter along chromosome or occurs more symmetrically on replicative strands. Given the importance of mutation signature analysis, it is therefore vital to control for mutation load when evaluating properties of signatures. By extension, other properties of mutational signatures such as nucleosome occupancy, transcription factor binding occupancy, or histone modifications may be subject to similar heterogeneity^10^.

Our controls for mutation numbers across mutation load groups, and the down-sampling of mutations in individual LCLs, indicate that the association between mutation load and the mutation landscape is not due to lack of statistical power. Instead, these appear to be two correlated attributes that are inherent to individual samples. We consider several possible mechanisms to explain this inter-sample variability. First, it is conceivable that past mutations inherently increase the probability, and skew the distribution of future mutations, in a type of mutational feedback loop. This could happen, for instance, due to local recruitment and retainment of mutagenic DNA repair pathways. However, the observation that SHM mutational clustering decreases with higher mutation load implies that mutation rate increases in late replicating regions are not driven by proximal changes, arguing against such a mechanism in LCLs. Instead, we favor a model by which the mutation of a *trans-*acting factor increases the global mutation rate and also underlies the shift of mutations towards later replicating genomic regions. As this mutation increases in clonal frequency, possibly due to compounding effects of the mutated gene(s) on cell proliferation, we would observe greater late replication timing bias for newly acquired somatic mutations. One candidate of interest we identified is *BCL6*, a cancer census gene prominently mutated in B cell lymphomas. *BCL6* is a transcription factor that prevents cell cycle arrest under the tremendous DNA damage of SHM^56^. Current models pose that *BCL6* mutations disrupt its negative regulation, promoting proliferation despite ongoing mutagenesis^56^. Further investigation on functional mutations of *BCL6* in B cells may elucidate its role in elevated late SHM replication timing bias with high mutation load. It would also be important to determine whether the mutation load effect is unique to SHM in B cell types, or if similar or other processes with comparable effects take place in other cell types. Regardless, we argue that mutation load, even if being a proxy for another underlying mutational landscape shift, is important to consider in any studies of mutational patterns.

Another unexpected finding of this work relates to the mutational landscape of the inactive X chromosome. We found that SHM was elevated on Xi in agreement with the chromosome’s late replication, while its mutations were unanticipatedly distributed with respect to the replication pattern of Xa. Furthermore, SHM showed elevated late Xa replication timing bias in high mutation load samples, as observed on autosomes. Clock-like mutations, on the other hand, were distributed with respect to the disordered replication of Xi. These findings were supported by male-female comparisons in CLL. These results suggest that replication timing may not directly modulate where SHM mutations occur. Instead, some yet unidentified correlated factor that is otherwise unaltered on Xi and serves as an epigenetic “memory” of its pre-inactivation state, may explain the landscape of SHM. Since gene expression, chromatin structure, and chromosome conformation are all effectively lost on the Xi alongside replication timing programming^70,71^, it is difficult for us to speculate on the nature of such a factor at this time.

A major and still not fully answered question in the human mutagenesis field pertains to the mechanisms that lead to preferential mutation accumulation in late replicating regions. The comparison of SHM and clock-like mutations on both the autosomes and the X-chromosome support the idea that there is no singular mechanism that can explain this association. Rather, mutational landscapes are shaped by composites of pathways with varied associations with the replication program. By first categorizing which pathways are directly modulated by replication timing, the underlying mechanisms may be more easily probed. Nevertheless, in combination with mutational pathways, mutational load, and rate of clustering, replication timing is an effective predictor and likely to be a critical driver of regional mutation rates across chromosomes. Given that replication timing itself is a polymorphic trait in humans^38,72^, we would predict that different people would have different mutational patterns in different genomic regions; characterizing such a form of genetic variation would require incorporating the multiple factors we described here, including mutational signature abundance, autosomal mutation load, and mutation clustering.

## Methods

### Genomic data sources and mutation calling

#### LCL genomic data sources

Mutations in the 1662 LCL offspring were sourced from six cohorts (**Table 1**). These offspring were matched to 989 pairs of fully genotyped parents, as 377 families contained two or more offspring. Eight families covered three generations. The largest cohort was iHART^73^ and included 1028 offspring with or without a diagnosis of autism. While iHART samples included both LCL and whole blood samples, only LCL offspring were included in this study, although for parental data we also considered whole blood samples (1.2% of parents). The second-largest LCL mutation cohort was sourced from the 1000 Genomes Project (1kGP) and contained 602 trios^74^. We used 49 offspring from the Polaris project Kids cohort^75^ as replicate samples as all overlapped the 1kGP cohort. An additional nine offspring were sourced from the Repeat Expansion (RE) cohort^76^ and included two fragile-X syndrome patients that we nonetheless have shown before do not have global replication timing alterations compared to healthy samples^77^. We sourced another 13 offspring from the Illumina Platinum^78^ family; of those, two (NA12878 and NA12877) overlapped with 1kGP samples and were used for primary analyses instead of the latter due to their higher read depth (∼50x compared to ∼30x).

We obtained 12 LCL trios from the Coriell Institute and sequenced and aligned them in-house. Samples were sequenced at Genewiz (South Plainfield, NJ) on Illumina HiSeq X (2×150bp) to a depth of approximately 15X (for further information, see Caballero et al. 2021^77^). Reads were converted into unaligned BAM files and marked for Illumina adaptors with Picard Tools (v1.138) (http://broadinstitute.github.io/picard/) commands ‘FastqToSam’ and ‘MarkIlluminaAdapters’. BAM files were then aligned to hg38 with BWA-mem^79^ (v0.7.17), and duplicate reads were marked with Picard Tools command ‘MarkDuplicates’. These alignment steps were similar to those implemented for the other LCL cohorts. Among these 12 offspring, two are affected by ataxia-telangiectasia yet did not show global replication timing alterations compared to healthy LCLs^77^.

#### LCL genotyping

In order to ultimately identify mutations, we first genotyped LCL offspring and parents. Genotypes for iHART samples were obtained from Ruzzo et al. 2019^73^. All other LCL cohorts were genotyped by us using the GATK (v4.1.4.0) best practices for germline short variant discovery^80,81^. Briefly, BAM files were recalibrated and aligned around common insertions and deletions with ‘BaseRecalibrator’ and ‘IndelRealigner’. Next, gVCF files were generated from all recalibrated BAM files using ‘HaplotypeCaller’. gVCFs were then merged into families with ‘CombineGVCFs’ and joint genotyped with ‘GenotypeGVCFs’. Finally, SNVs were recalibrated with ‘VariantRecalibrator’. We note that genotype calling for the iHART cohort differed from the above in that all samples were jointly genotyped, and variants were removed if they had a depth of <10X, a genotype quality of <25, or an alternative allele frequency of <0.2; we subsequently applied equal or stricter filtering metrics to all samples when identifying mutations, hence ruling our an effect of these differences in iHART genotyping on our analyses.

For samples originally aligned and genotyped in hg19 (approximately half of all samples), genotypes were lifted-over to hg38 coordinates using vcf-liftover (https://github.com/hmgu-itg/VCF-liftover, only liftover within the same chromosome were allowed). We removed genotypes in samples originally aligned to hg38 at coordinates without an hg19 equivalent to compensate for the reduction of genotypes following liftover. This eliminated approximately 1.9% of all sites.

#### LCL mutation calling

Candidate mutations were identified as single nucleotide Mendelian errors between parent and offspring alleles. The following steps were based on previously established family-based mutation calling methods from Yuen et al. 2016^82^. Mutations on the autosomes and X-chromosome in female offspring were identified as heterozygous genotypes (for the reference allele and an alternate allele) in offspring where parents were homozygous for the reference allele. For the X-chromosome in male offspring, mutations were identified as sites with only an alternate allele where the mother is homozygous for the reference allele. Next, we filtered mutations with a Fisher’s exact test Phred-scaled p-value (FS)<60.0, RMS mapping quality (MQ)< 0.0, Wilcoxon rank sum test z-score of mapping qualities (MQRankSum)<-12.5 or read position (RPRS)<-8.0, symmetric odds ratio (SOR)>3, and a Phred-scaled quality score (QUAL)<30. We excluded sites that did not pass variant quality score recalibration. To remove sub-clonal mutations and potential technical errors, we eliminated candidate mutations for which the mutant (alternate) allele frequency was <0.2. We removed likely inherited variants where either parent contained reads matching the mutant allele. Finally, to eliminate possible false-positive mutation calls caused by somatic deletions in the offspring (and hence reduced genotyping accuracy), we eliminated candidate mutations in cases where the offspring read depth was <10% of the combined parental read depth (again, adjusted for the X-chromosome in male offspring) at the mutation site. After this initial hard filtering, 4.4 million candidate mutations were called across all 1662 offspring.

Next, we removed candidate mutations based on genomic location. We first removed 61,479 candidate mutations around the HLA locus (chr6:28477797–33548354 in hg38) due to the high propensity for genotyping errors stemming from high local polymorphism density^83^. Similarly, we removed 63,547 mutations around the immunoglobulin heavy locus (*IGHV*, chr14:105580000-106880000 in hg38), which is hypermutated in LCLs. Next, we removed 587,511 mutations within gaps >25Kb in the LCL replication timing profile (see section **Replication timing profiles**). Regions of the genome removed for HLA and *IGHV* were also removed from the LCL reference RT profile.

To further eliminate inherited variants, we implemented a last filtering step to remove mutations based on population allele frequency. Specifically, we removed mutations with a gnomAD^84^ V3 allele frequency of >0.001. We did not use a frequency of zero as many of our samples (including all 1kGP individuals), and their somatic mutations, are represented in gnomAD. We also filtered mutations occurring in more than 30 of the 1662 offspring. In total, 2,826,985 candidate mutations were eliminated through this allele frequency filtering. After all filtering steps, 885,655 autosomal and 42,061 X-chromosome mutations remained in the 1662 non-replicate LCL offspring.

For each mutation, trinucleotide context was generated with SigProfilerMatrixGenerator^85^, and replication timing values at mutations sites were calculated with the R function ‘approx’ using the linear method.

#### LCL mutation validation

Parent-offspring mutation calling carries a risk of falsely identifying an inherited variant as a *de novo* mutation. This could stem, for instance, from failing to identify the inherited alleles in a parent due to a somatic deletion or false-negative genotyping. To quantify the proportion of false mutations that are inherited variants, we analyzed mutation calls in 73 monozygotic (MZ) twin pairs. MZ twins share all inherited alleles and germline mutations but have unique somatic mutations (**Fig S1B**). Although parent-offspring mutation calling cannot distinguish somatic from germline mutations, having an estimate for one of those enables to estimate the other. Specifically, based on all samples from denovo-db^86^, the average human contains 65.5 autosomal germline mutations. In contrast, in this study, MZ pairs shared between 81 and 245 autosomal mutations (median:113; **Fig S1C, D**). Thus, the excess number (above 65.5) of MZ twin shared mutations provides a rough estimate of the number of falsely called mutations that are likely inherited variants (**Fig S1E**). We thus predicted that between 1.85% to 27.2% of autosomal mutations in MZ twins are inherited variants (median: 9.66%; **Fig S1E**). This is likely an overestimate, as the paternal age among MZ twins was relatively high (median: 32.26 years, range: 20.43-78.51), thus increasing the expected number of germline mutations.

We also estimated false mutation calls derived from technical errors by analyzing genotype calls in 51 offspring that were resequenced by different groups on different platforms (**Table S1**). We compared mutant alleles of samples in the main dataset to the GVCF of the replicate. A mutation was considered validated if the mutant allele was found in the replicate sample at any frequency. A median of 93.1% of autosomal mutations were supported by their replicate sample (range: 65.1-98.7%; **Fig S1F**). The mutations that could not be validated did not show a strong enrichment towards late replication timing and, therefore, should not have influenced our results (**Fig S1G**). We further validated mutation calls in the offspring sample NA12878. The Illumina Platinum cohort sample of NA12878 was used as part of the main dataset (of 1662 offspring), and the 1kGP NA12878 sample was used for validation (and counted as part of the 51 replicate sample analysis mentioned above). We sourced four other replicate sequencings of NA12878 (**Table S1**) and found that 98.8% of mutations were supported by at least one alternate source.

#### CLL mutation data

Mutations in chronic lymphocytic leukemia (CLL) patients were obtained from the ICGC/PCAWG cohorts CLLE-ES. Alignment and mutation calling for tumor samples (peripheral blood-derived) and normal samples was performed by PCAWG using their pipeline^87^ in hg19. We only included mutations called from 151 patients with whole genome sequencing. This provided 371,252 autosomal mutations and 23,130 X-chromosome mutations.

Before filtering, all mutations were lifted to hg38 using the vcf-liftover method, as used in LCL. We then removed mutations around the HLA and IGHV loci and in gaps of the LCL replication timing profile. Hence, we used two LCL replication timing profiles in our analyses: one in which regions filtered from the LCL offspring dataset were removed, and another in which regions filtered from the CLL dataset were removed. We interpolated replication timing values for the final 355,474 autosomal and 22,131 X-chromosome mutations with the CLL-filtered LCL reference replication timing profile and determined trinucleotide contexts in an identical manner to LCLs.

#### HCT116, HT115, and LS180 mutation data

The HCT116 line was a gift from the tissue culture lab at the Francis Crick Institute. Cells were grown in Dulbecco’s Modified Eagle Medium (DMEM), 10% fetal calf serum, penicillin, and streptomycin. Culture was maintained at 37°C with 5% CO2. Passage was performed approximately twice per week for one year. BAM files were generated by aligning reads to hg38 and recalibrated in an identical manner to our processing of the LCL data as described above. BAM files from the passage of HT115 and LS180 were sourced from Petljak et al. 2019^25^. BAM files, originally generated by aligning reads to hg19, were recalibrated identically to our processing of LCL data (above). For LS180 and HT115, we lifted mutations to hg38 (as described above).

Mutations in HCT116 were identified with GATK (v4.1.4.0) mutect2^88^ per the somatic short variant discovery best-practices pipeline. The parental clone was considered the normal sample, and daughter clones were considered tumor samples. For filtering, read orientation bias artifacts were predicted with the command ‘LearnReadOrientationModel’ and used in filtering with ‘FilterMutectCalls.’ The Mutect2 step of cross-sample contamination was not implemented since the samples were cell lines. We identified candidate mutations as heterozygous calls that passed the mutect2 filtering and were unique to a daughter subclone. We required that at daughter candidate mutation sites, the parental genotype must be homozygous for the reference allele and not contain any mutant allele reads. We removed mutations where the parental clone had no read depth, as this prevented confident mutation calling. Finally, we only retained candidate mutations with an MQ of <40 and an alternate (mutant) allele frequency of >0.2 and <0.8 in the daughter.

We removed mutations in all colon adenocarcinoma cell lines around the HLA locus and gaps >25Kb in the respective cell type replication timing profile. The final mutation dataset contained 150,470 autosomal mutations in the six HCT116 subclones, 28,944 autosomal mutations in the five HT115 subclones, and 14,974 autosomal mutations in the five LS180 subclones. Mutation trinucleotide context and interpolated replication timing values were assigned using the methods described above for LCLs and CLL.

### Replication timing profiles

#### LCL

The LCL replication profile was generated using TIGER^37^ from median read count data from all 1662 offspring. First, uniquely mapping reads were extracted from aligned BAM files of each sample. For samples aligned to hg19, BAM coordinates were lifted to hg38 in an identical manner to mutations. We compensated for lift-over by modifying TIGER to exclude hg38 coordinates with no hg19 equivalent when creating 2.5Kb windows of uniquely alignable sequence. We tested the effect of this method by comparing the replication timing profiles of 22 samples originally aligned to hg38 with those aligned to hg19 and lifted-over to hg38. The lifted replication timing profile in all samples on all autosomes was nearly identical (Pearson’s *r* >0.99) to the one aligned to hg38.

Using default TIGER parameters, the liftover-corrected 2.5Kb windows were GC-corrected and normalized to an autosomal genome copy number of two. We eliminated subclonal aneuploidies in individual offspring by filtering out whole chromosomes with an average autosomal copy number of >2.2 or <1.8, an X-chromosome copy number of >2 or <1.6 for female offspring, and an X-chromosome copy number of >1.2 or <0.8 for male offspring. This removed 34 chromosomes in 23 samples. We removed suspected small copy number alterations by filtering out 2.5Kb windows with an exceptionally high or low median copy number across all offspring and within individual offspring. We first removed autosomal and female X-chromosome windows across all offspring with a median copy number ±0.6 than that chromosome’s median copy number (as calculated from all offspring). The cutoff was ±0.4 for the X-chromosome in male offspring. We then filtered out windows in individual offspring with a copy number ±0.6 than that chromosome’s median copy number (as calculated in the individual offspring). The cutoff was ±0.3 for the X-chromosome in male offspring. We next calculated autocorrelation for all offspring using the MATLAB command “autocorr” and removed whole chromosomes for samples with abnormally high autocorrelation. This removed 51 chromosomes in 26 samples. Finally, we discarded the two offspring, HG02523 and NA12344, as they had more than six individual chromosomes removed.

Mutations in LCL offspring and HCT116 daughter subclones were not removed if an offspring’s chromosome was filtered out during replication timing generation. However, as previously mentioned, candidate mutations were removed in regions >25Kb where replication timing was not available for all offspring. This arose from windows filtered out for disproportionately high or low median copy number across all offspring, which removed 92Mb on autosomes (3.67% of the autosomal genome).

After filtering, we took the median GC-corrected data in 2.5Kb each window across all offspring. For the X-chromosome, we calculated separate medians using only male or female offspring. Replication timing values were generated by smoothing the median GC-corrected data with a cubic smoothing spline (MATLAB command ‘csaps’, smoothing parameter: 1×10^−17^). Only regions of >20 continuous 2500bp windows were included. Smoothing was not performed over data gaps >100Kb or reference genome gaps >50Kb. The smoothed profiles were then normalized to an autosomal mean of zero and a standard deviation of one. For analyses on the X-chromosome, we generated an X-chromosome replication timing profile considering only male LCL offspring.

We compared our median LCL replication timing profile to a replication profile of NA12878 generated by sequencing S and G1 phase DNA^89^. The S/G1 coordinates were interpolated to TIGER window coordinates with the MATLAB function ‘interp1’. The LCL replication timing used in this study highly correlated to the S/G1 profile (Pearson’s *r* = 0.94; **Fig S1J**).

#### HCT116

We similarly generated a median autosomal replication timing profile for HCT116 from the six daughter subclones and the parental line using TIGER. Liftover adjustment was not implemented as all samples were originally aligned to hg38. HCT116 is nearly diploid, with several large copy number alterations present in some or all samples. As in LCL, we removed these copy number alterations by filtering out 2.5Kb windows in individual samples with a copy number ±0.6 than the chromosomal median copy number (as calculated in the individual sample). Each sample was then filtered via the TIGER command ‘TIGER_segment_filt’ (using the MATLAB function ‘segment’, R2: 0.04, standard deviation threshold: 2.5). After filtering, we took the median GC-corrected data in 2.5Kb each window across all samples. Altogether, 280Mb were removed in filtering (11.1% of the autosomal genome). Notably, four copy number alterations >10Mb were removed from all samples.

#### HT115 and LS180

HT115 and LS180 replication timing profiles were generated from S/G1 sequencing as described in Massey et al., 2019^89^. DNA from each cell cycle fraction was sequenced using an Illumina NextSeq 500 and aligned to hg19. The S/G1 DNA replication timing profile for HT115 was previously described^21^. The S/G1 replication timing coordinates were lifted to hg38 as described above for LCLs.

We compared the final TIGER-generated HCT116 replication timing profile to one generated by S/G1 alongside HT115 and LS180. The two profiles were highly correlated (Pearson’s *r* = 0.91; **Fig S1J**). We chose to use the TIGER-generated profile for HCT116 to match the source of the mutation calls.

### Mutation counts and signature fitting

We fit the previously described biologically relevant COSMIC v3.2 SBS signatures^1^ to all autosomal mutations in the five cell types using the MutationalPatterns^90^ command ‘fit_to_signatures‘. Following current best-practices^45^, individual COSMIC signatures were corrected by adjusting the 96 trinucleotide frequencies by the relative abundance of trinucleotide frequencies between the filtered and unfiltered autosomal genome. We used cosine similarity to assess the confidence of signature fit. This metric compares the original trinucleotide frequencies of mutations to reconstructed frequencies based on predicted signature contributions. A value of one indicates an identical reconstruction. We calculated cosine similarity with the MutationalPatterns command ‘cos_sim’. We additionally performed 1000 bootstrap sampling when fitting signatures using the MutationalPatterns command ‘fit_to_signatures_bootstrapped’. We used the standard deviation of 1000 bootstrap samples as the standard error for signature contribution. Standard errors for combined signatures (e.g., MMRd, which is the combination of SBS21 and SBS44 in HCT116/LS180) were calculated using standard error in the difference of the means (the square-root of the sum of variances).

To assess the relationship of mutations or signature abundance to replication timing, we divided the autosomal replication timing profiles of each cell type into 20 bins ordered by replication timing. Each bin contained an equal 5% of the genome. In later analyses where mutations were reduced (e.g., stratification by replicative strand), we used five bins (each with an equal 20%) to preserve resolution. The number of bins was chosen to optimize visualization for the different analyses. When fitting signatures to mutations, we again corrected for trinucleotide abundances within each replication timing bin. For this, the 96 trinucleotide frequencies were corrected by the relative abundance of trinucleotide frequencies between the filtered and unfiltered autosomal genome within the replication timing range of each bin.

### Replicative strand asymmetry

The local slope of replication timing provides replicative strand information for the positive strand of the genome. We assigned 2.5Kb smoothed data windows of positive slope (based on the immediate flanking windows) as lagging replicative strand on the positive genome strand and leading replicative strand on the negative genome strand. Reciprocally, windows of negative slope were assigned as leading replicative strand on the positive strand and lagging replicative strand on the negative strand. At locations of a slope change, flanking windows within 100Kb were assigned undefined replicative strandedness for both the positive and negative genome strands. Undefined replicative strandedness comprised 600.15Mb (approximately 25%) of the LCL replication timing profile, 599.49Mb in CLL, 740.15Mb in HCT116, 1113.77Mb in LS180, and 1000.07Mb in HT115. Mutations were partitioned into leading or lagging groups based on (1) whether the pyrimidine base of the substitution was on the positive or negative genome strand and (2) the replicative strand of the positive and negative genome strands at that coordinate. We did not include mutations in regions of undefined replicative strand in asymmetry analysis.

We fit the biologically relevant mutational signatures separately to replicative strand-partitioned autosomal mutations. As performed above, individual COSMIC signatures were corrected by adjusting the 96 trinucleotide frequencies by the relative abundance of trinucleotide frequencies between the filtered leading or lagging replicative strand and unfiltered autosomal genome. Regions of undefined strandedness were not included in correction. To assess the relationship of mutational replicative strand asymmetry to replication timing, we divided the autosomal replication timing profile (voiding regions of undefined strandedness) into five bins ordered by replication timing value. Each bin contained an equal quintile (20%) of the genome. We fit the biologically relevant mutational signatures separately to the replicative strand-partitioned mutations in each quintile. Again, we performed signature correction using only regions of defined strandedness within the range of replication timing quintiles.

Before determining asymmetry values, we calculated replicative strand ratios for a given mutational signature using the formula:

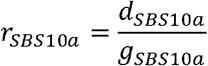

where *d* and *g* represent the number of autosomal mutations on the respective leading and lagging strand regarding the genomic strand of the substituted pyrimidine base.

As described above, we calculated standard error for a signature as the standard deviation of 1000 bootstrap samples. Standard error was calculated separately for mutations partitioned to the leading and lagging replicative strand. To get standard error for a replicative strand ratio, we propagated standard errors from the leading and lagging strands using the formula:

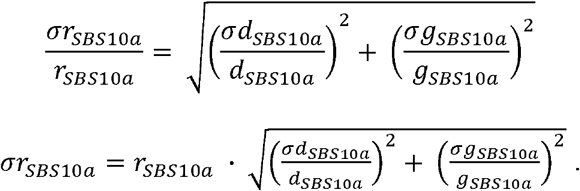

We then calculated replicative strand asymmetry values using the formula:

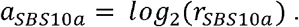

To calculate standard error for asymmetry values, we subtracted the error from the replicative strand ratio before log2 transformation. Thus, we determined the error for asymmetry as:

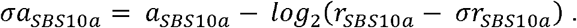

To increase strand asymmetry confidence, we repeated the analysis of strand asymmetry in LCL, CLL, and HCT116 while removing 500Kb (instead of 100Kb) around regions of slope change. The rationale for this validation was that origin and termination sites in replication timing profiles may be regionally imprecise or variable across samples, leading to false mutation strand assignment even after removing 200Kb around regions of slope change. HT115 and LS180 were not included in this reanalysis due to an insufficient number of mutations.

### Gene associations for late replication timing bias

We identified individual LCL mutational replication timing bias by calculating the proportion of mutations in four replication timing bins. We used the linear slope of proportions as a representation for replication timing bias and calculated PCs using the R command ‘prcomp.’ Gene associations were calculated using the binary state of whether at least one mutation fell within the range of a protein coding gene (https://ftp.ncbi.nlm.nih.gov/genomes/all/GCF/000/001/405/GCF_000001405.39_GRCh38.p13/) against individual replication timing biases. Mutation functionality was not considered. P-value of association was calculated with the R command ‘lm’ and individual autosomal mutation load was inputted as a covariate. 97 genes showed significant association for late replication timing biases and were mutated in at least 50 samples.

### Clustering mutations

We clustered SHM-context mutations, which represented 26.69% of autosomal LCL mutations and 21.13% of CLL mutations, using ‘ClusteredMutations’ (https://cran.r-project.org/web/packages/ClusteredMutations/index.html) command ‘showers.’ The minimum cluster size was two mutations, and the maximum distance between SHM-context mutations was 500bp. We simulated autosomal SHM-context mutations of matched mutation rates in 20 replication timing bins. Within the replication timing range of each bin, we performed 1000 random selections of SHM-context motifs (TA, TT, or AA loci on the positive genome strand) without replacement. The simulated mutations were clustered identically as described above for real mutations.

We evaluated the distance of SHM-context mutations to 22,337 protein-coding genes and the C>N mutations in LCL offspring and CLL. We defined genes as all transcribed sequences (mRNA in the gene feature table), including introns and UTRs. As many gene models overlapped, we merged intervals using the bedtools^91^ (v2.29.2) command ‘merge.’ We interpolated LCL replication timing values using the center coordinate of the merged gene regions. We calculated the distance between SHM-context mutations and gene/C>N mutations with the bed tools command ‘closest.’

### Determining Xi parental identity and phasing mutations

We phased Mendelian inherited single nucleotide variants in female LCL offspring. For each variant, we required the offspring and parents to have a read depth ≥5, MQ>30, FS<60.0, MQRankSum>-12.5, RPRS>-8.0, and SOR<3. In the heterozygous offspring genotype, we required the alternate allele frequency to be greater than 0.3. We calculated parental copy number disparity as the absolute difference of mean sequencing read depth for paternal and maternal alleles divided by their combined read depth. To determine a threshold for identifying X-inactivation, we used the 95^th^ percentile of parental copy number disparity on chromosome 14. This chromosome was chosen as it contained the most comparable number of phaseable variants as chromosome X. The parental identity of Xi was assigned to the parental homolog with the lower mean sequencing read depth.

We phased mutations occurring on the same read or mate-pair as a phaseable inherited variant. We first determined the read names containing the maternal and paternal alleles using the Samtools^92^ (v1.6) command ‘mpileup.’ We repeated this process to identify read names containing the mutation alleles. We phased mutations where read names containing mutation alleles exclusively matched those phased to one parent. If mutation alleles matched read names phased to both parents, the mutation was considered ambiguous. We calculated mutational signature contributions on phased chromosomes as described above using the biologically relevant LCL signatures corrected for individual chromosome trinucleotide content.

## Data and code availability

All replication timing profiles in hg38 coordinates and relevant code are available in the supplementary information. BAM files for HCT116 and relevant S/G1 profiles are available as SRA bioproject PRJNA875498. Mutation counts for LCL offspring, CLL-M/U predictions, and Xi parental identity predictions are available in Table S1.

## Acknowledgements

We thank Verena Höfer for technical assistance in generating data for HCT116. This work was funded by the National Institutes of Health (award DP2-GM123495 to AK), the National Science Foundation (award MCB-1921341 to AK), and the United States-Israel Binational Science Foundation (award 202108 to AK and I. Simon).

