## Supplemental Figures for "Multifactorial heterogeneity of the human mutation landscape related to DNA replication dynamics"

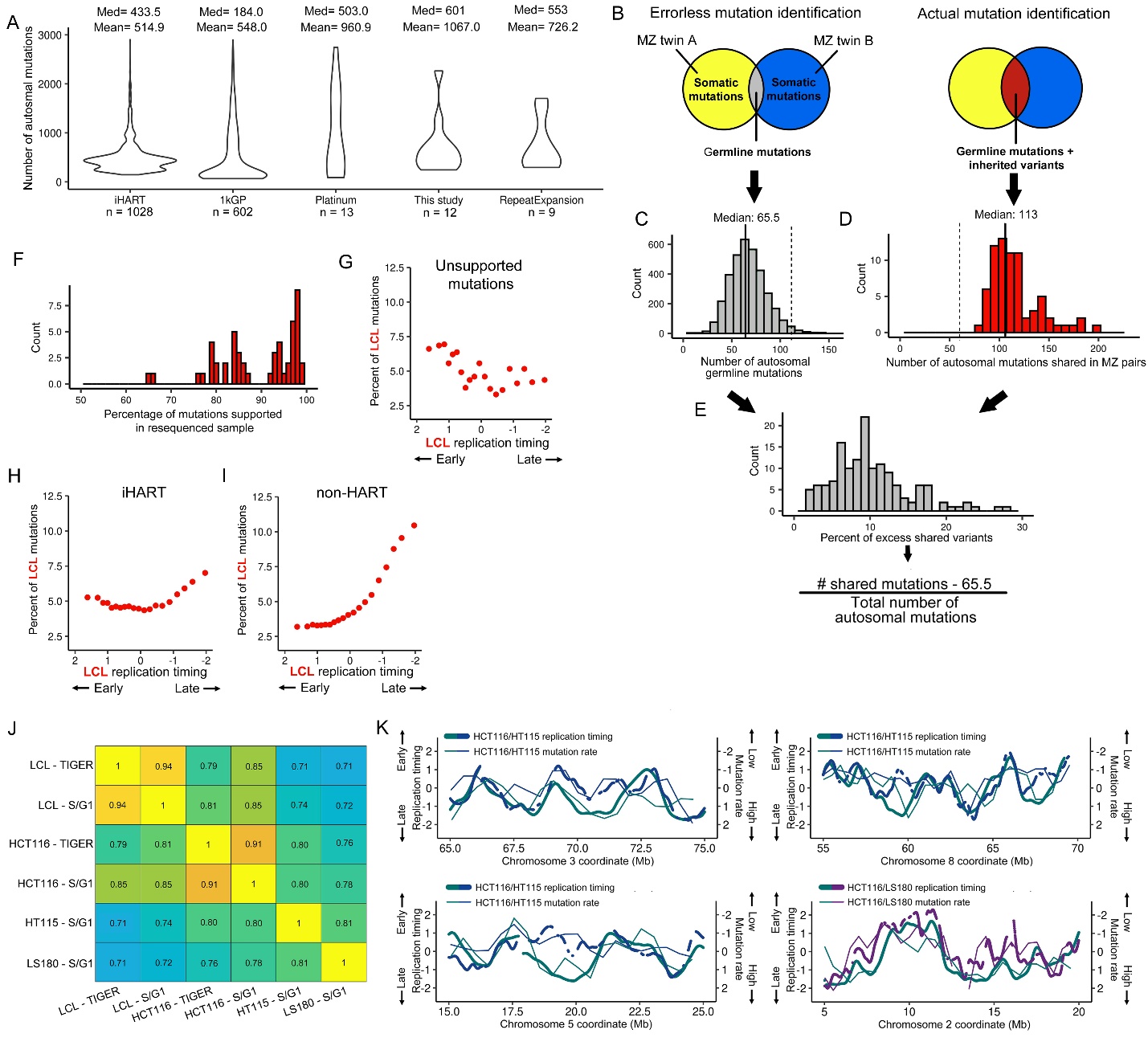

**Fig S1. Quality control of mutations called in LCLs and comparison of replication timing profiles and mutation rate correlations in the different cell types/lines.** (A) The number of mutations identified across all autosomes in each sequencing cohort. (B-E) Using monozygotic twin (MZ) pairs to identify the proportion of inherited variants. (B) Venn diagrams demonstrate the over-representation of shared mutations caused by inherited variants where parental alleles were not identified. (C) The number of germline mutations per individual from denovo-db (denovo-db.gs.washington.edu). We considered this the expected number of mutations shared in MZ twins. (D) The actual distribution of the number of shared mutations in LCL MZ pairs. (E) The fraction of mutations estimated to be inherited variants. The number of shared mutations minus the expected number of germline mutations was divided by the total number of mutations for each MZ sample. (F) The percentage of mutations supported in a single replicate sequencing across 51 LCL offspring. (G) The distribution of mutation counts in 20 replication timing bins for mutations not supported in the 51 LCL samples sequenced in replicate. (H) Distribution of LCL mutations from the iHART cohort. (I) Distribution of LCL mutations from samples not in the iHART cohort**.** (J) Pearson correlations of autosomal replication timing profiles. (K) As in Fig 1K, mutation rate correlates to the cell type-specific replication timing in HCT116, LS180, and HT115.

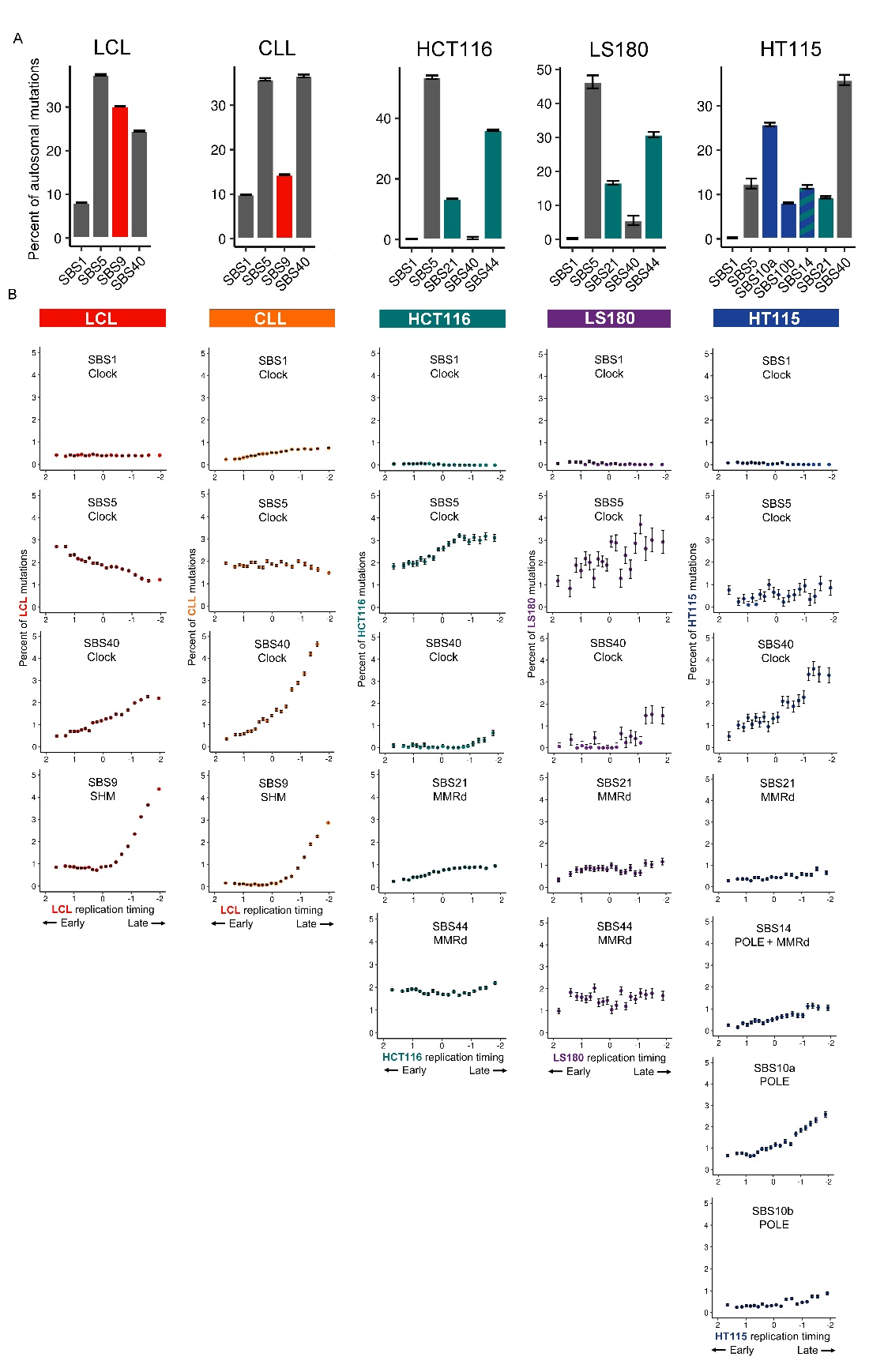

**Fig S2 Mutational signature abundance as a function of replication timing in all cell types/lines.** (A) Bar plot representation of Fig 2A-E. (B) The proportion of autosomal mutations contributed by various signatures in 20 replication timing bins of uniform genome content. For all panels, error bars represent the standard error of signature fit (± the standard deviation of 1000 bootstrapped samples).

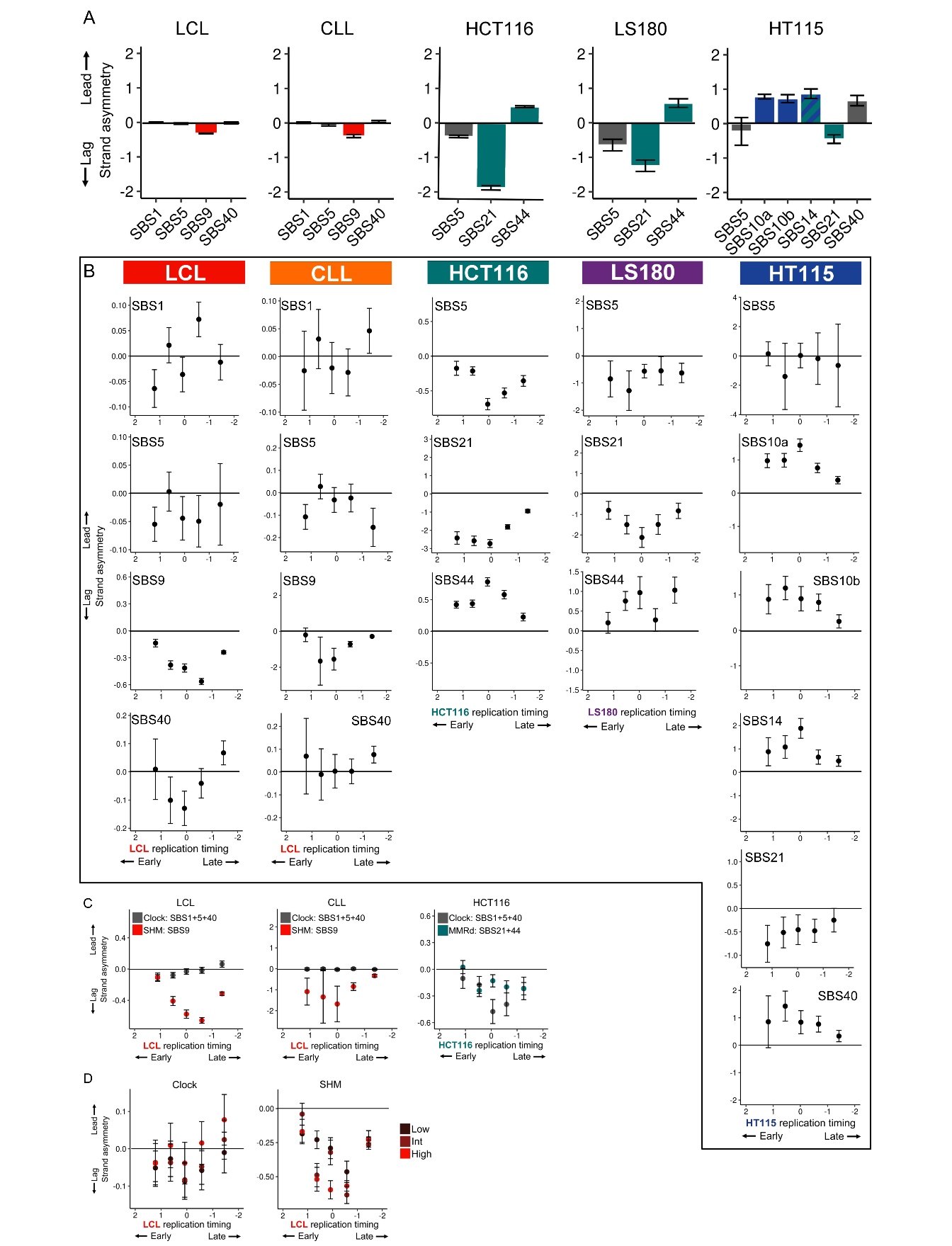

**Fig S3.** **Replicative strand asymmetry for mutational signatures and mutation load groups.** (A) Mutational strand asymmetry for signatures comprising at least 5% of autosomal mutations. (B) Replicative strand asymmetry for mutational signatures in five replication timing bins of uniform genome content. Error bars for all panels represent the standard error of replicative asymmetry. (C) As in Fig 3C, I, K, replicative strand asymmetry for mutational categories in five replication timing bins of uniform genome content with 500Kb regions removed flanking slope directionality changes. (D) Extended from Fig 3E, G, clock-like and SHM mutational asymmetry in all LCL mutation load groups as a function of replication timing. Error bars for all panels represent the standard error of replicative asymmetry.

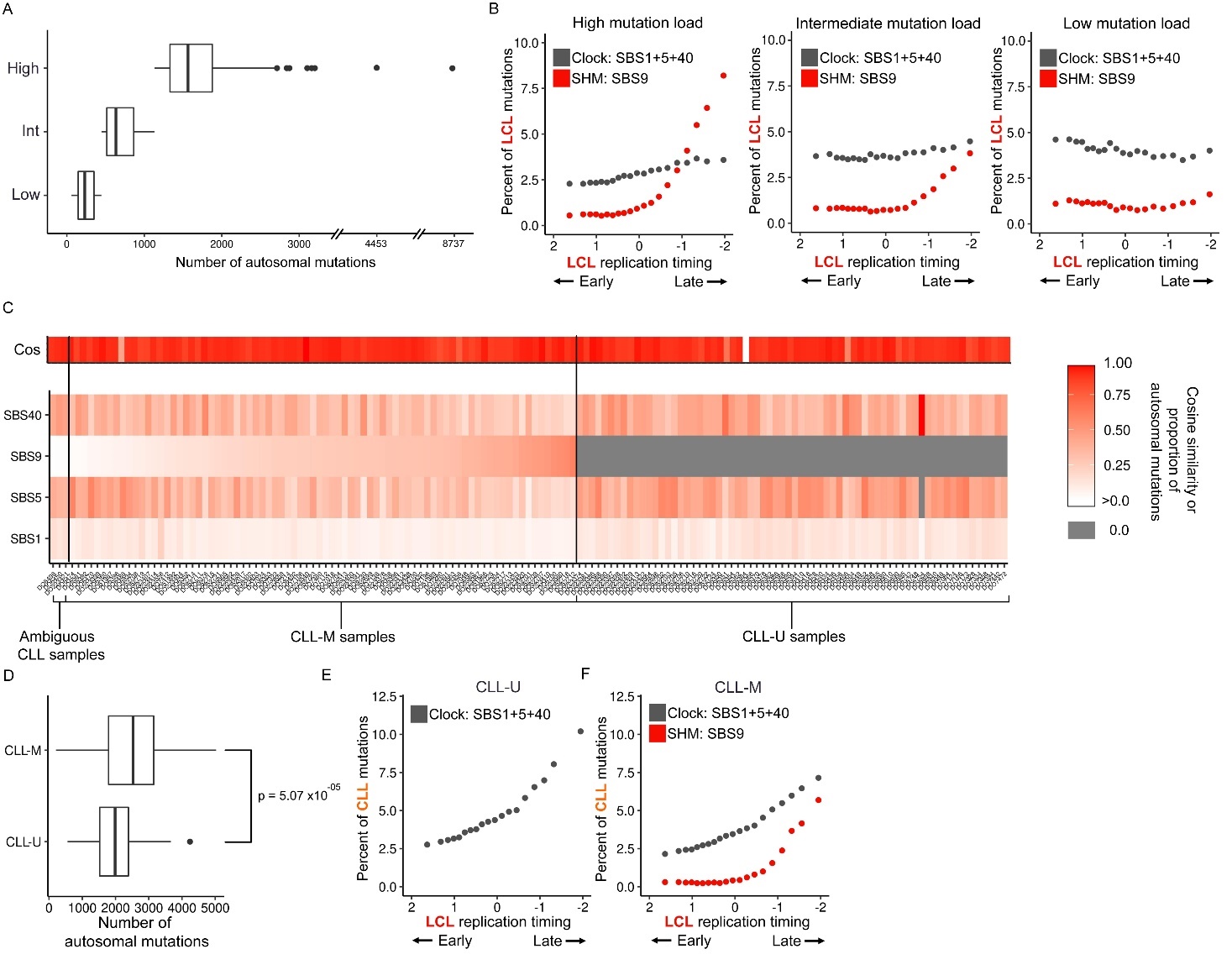

**Fig S4**. **Stratification of LCLs by autosomal mutation load and CLL samples by IGHV mutation status.** (A) The distribution of total autosomal mutations in LCL offspring in the low, intermediate, and high mutation load groups. (B) Clock-like and SHM signature abundance as a function of replication timing in the LCL mutation load groups. (C) The abundance and cosine similarity of mutational signatures fit to individual CLL samples. (D) The distribution of total autosomal mutations in CLL-M and CLL-U samples. (F) Clock-like and SHM abundance as a function of replication timing in the CLL-U and CLL-M samples.

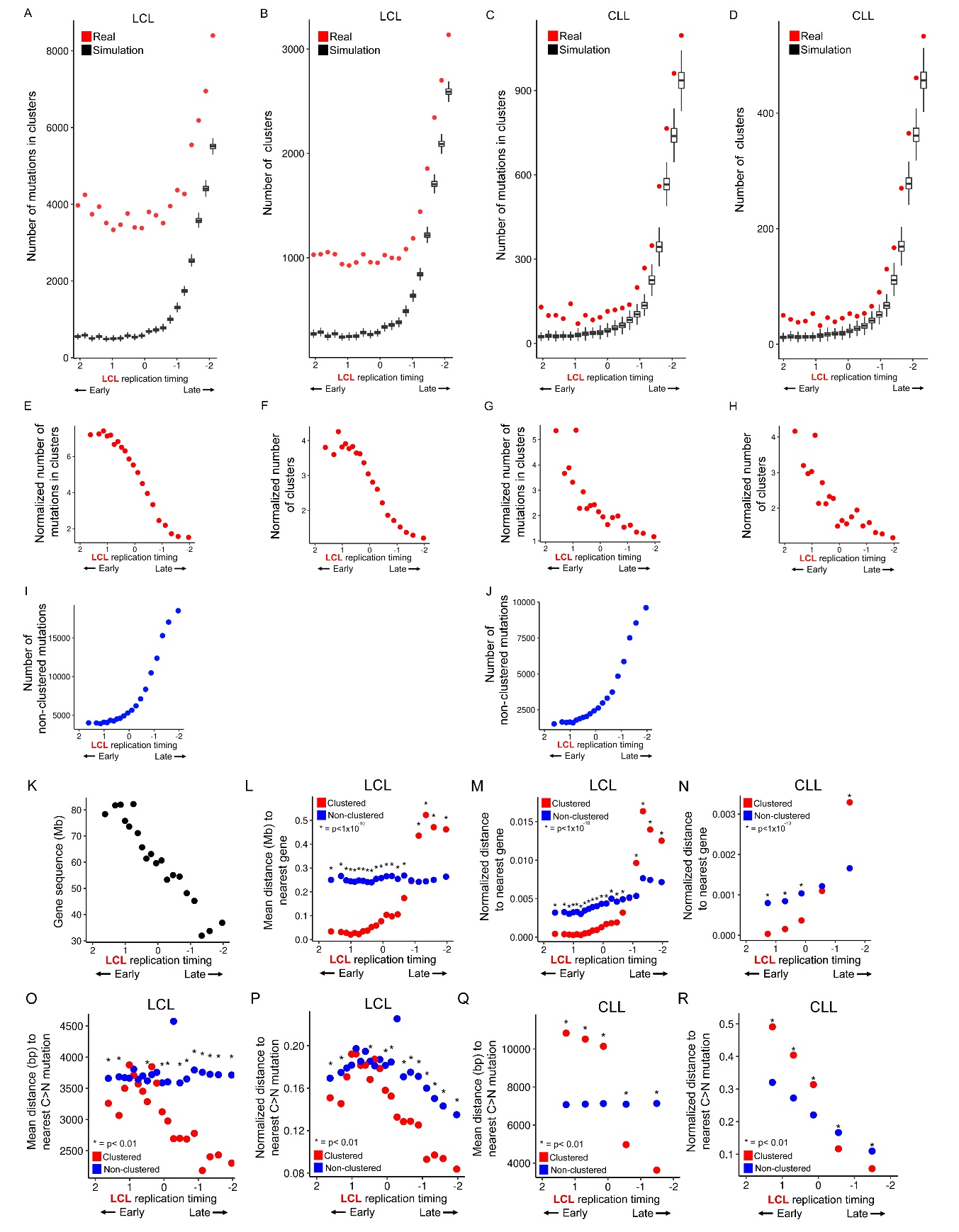

**Fig S5. Clustering of SHM-context mutations.** (A-B) The number of LCL SHM-context mutations within clusters (A) and the number of clusters (B) compared to 1000 iterations of simulated and clustered SHM-context mutations. Real and simulated clustering rates are within 20 replication timing bins of uniform genome content. (C-D) As in panels A and B for CLL SHM-context mutations. (E-F) The number of LCL SHM-context mutations within clusters (E) and the number of clusters (F) normalized by the simulated mean within replication timing bins. (G-H) As in panels E and F for CLL SHM-context mutations. (I-J) The number of non-clustered LCL (I) and CLL (J) SHM-context mutations within replication timing bins. (K) The total sequence length of protein coding genes in 20 replication timing bins of uniform genome content. (L) The mean distance of LCL clustered/non-clustered mutations to the nearest gene in 20 replication timing bins. (M) The data presented in panel L normalized by the length of protein coding sequence genes in panel K. (N) As in panel K for CLL in five replication timing bins of uniform genome content. (O) The mean distance of LCL clustered/non-clustered mutations to the nearest C>N mutation in 20 replication timing bins. (P) The data presented in panel O normalized by C>N mutation count per bin. (Q-R) As in panels O-P, for CLL. The mean distance to nearest C>N mutation (Q), and distance normalized by C>N mutation count (R).

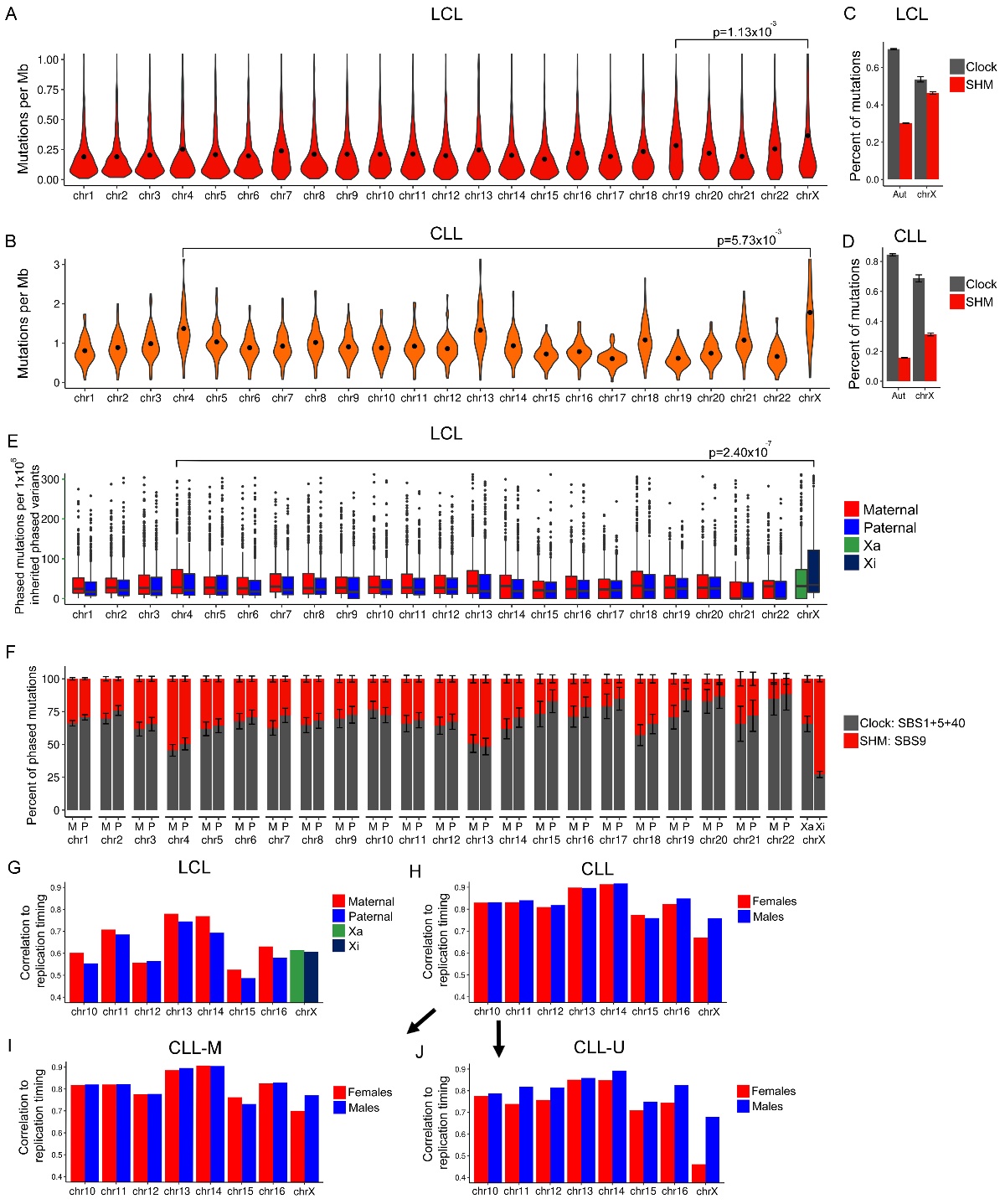

**Fig S6**. **Mutation rates across chromosomes using unphased and phased mutations.** (A) Number of mutations in the 746 female LCL offspring normalized by chromosome length. The significance for mutation rate difference between chromosomes X and 19 is highlighted as it represents the most significant p-value. (B) As in panel A for the 59 female CLL patients. (C) Mutation pathway abundances in the female LCL offspring across all autosomes and the X-chromosome. In panels C-D and F, error bars represent the standard error of signature fit. (D) As in panel C for the CLL female patients. (E) Number of phased mutations normalized by the number of inherited phaseable variants per chromosome. The significance for mutation rate between Xi and maternally-phased mutations on chromosome 4 represents the most significant p-value. (F) Mutation pathway abundances for phased mutations. (G) Pearson’s correlation coefficient of regional phased mutation rates and replication timing. For Xa and Xi mutations, correlation was calculated using male X-chromosome replication timing. Regional mutation rates were calculated as the mean number of phased mutations across all samples in a 1Mb sliding window with a 0.5Mb step. (H-J) Pearson’s correlation coefficient of male and female CLL (H), CLL-M (I), and CLL-U (J) mutation rates and replication timing. For X-chromosome, correlation was calculated using male LCL X-chromosome replication timing. Regional mutation rates were calculated as in panel G.
