## Supplementary material for "Multifactorial heterogeneity of the human mutation landscape related to DNA replication dynamics": cluster_mutations.R

```
#######################################################
# Caballero et al. 2022
# Description: Clustering SHM context mutations. 
#  Performed for whole genome.
#######################################################

# Note, similar steps for replication timing bins. You just have to
# partition "SHM_context_mutations.csv" into bins. 

## INPUTS:
MUTATIONS = read.csv(file = "SHM_context_mutations.csv", header = FALSE)

## LIB:
library(ClusteredMutations)
library(seriation)

system("rm clusters.csv")
system("rm cluster_mutations.csv")

mut_in_clusters = 0
num_clusters = 0
num_mut = 0

for(i in 1:22){
  match = paste0("chr", i)
  coordinates = MUTATIONS$V2[MUTATIONS$V1 == match]
  num_mut = num_mut + length(coordinates)

  clusters <- showers(data = NULL, chr = NULL, position = as.numeric(coordinates), min = 2, max = 500)		
  mut_in_clusters = mut_in_clusters + sum(clusters$number) 
  num_clusters = num_clusters + length(clusters$number)
  clusters$chr = i
  clusters = as.matrix(clusters)
  for (clust in 1:length(clusters[,1])){
    sub = (MUTATIONS$V1 == match) & (MUTATIONS$V2 >= clusters[clust,3]) & (MUTATIONS$V2 <= clusters[clust,2])
    mut_info = as.matrix(cbind(IN$V1[sub], MUTATIONS$V2[sub],  MUTATIONS$V3[sub], MUTATIONS$V4[sub], MUTATIONS$V5[sub], MUTATIONS$V8[sub], MUTATIONS$V11[sub]))
    RT = MUTATIONS$V8[sub]
    cat(clusters[clust,], RT[1], "\n", sep = ",", file = paste0("clusters.csv"), append = TRUE)
    for (j in 1:length(mut_info[,1])){
      cat(mut_info[j,], "\n", sep = ",", file = paste0("cluster_mutations.csv"), append = TRUE)
    }
  }
}

cat("Num mut in clusters:", "\t", mut_in_clusters, "\t", 100*(mut_in_clusters / num_mut), "%\n")
cat("Num clusters:", "\t", num_clusters, "\n")	

# Non-clustering mutations can then be extracted from the original SHM context mutation file
```
