## Supplementary material for "Multifactorial heterogeneity of the human mutation landscape related to DNA replication dynamics": correct_signatures_whole_genome.pl

```
#######################################################
# Caballero et al. 2022
# Description: Correcting signature trinucleotides
#  for autosomes and the regions of the genome
#  used in this analysis (known replication timing)
#######################################################

# Note, for correcting in bins of replication timing, this
# is the same approach. The correction is the nucleotide
# content in the replication timing bin compared to the
# whole reference genome. 

#!usr/bin/perl
no warnings;
use Bio::Index::Fasta;
use Bio::Seq;

open(RT,"/path/to/replication/timing/csv");

$file_name = "/path/to/reference/genome/GRCh38_full_analysis_set_plus_decoy_hla.fa" . ".idx";                                          
$inx = Bio::Index::Fasta->new(-filename => $file_name, -write_flag => 1);
$inx->make_index("/path/to/reference/genome/GRCh38_full_analysis_set_plus_decoy_hla.fa");
$inx = Bio::Index::Fasta->new(-filename => $file_name);

# Fetching nucleotides in the genome with defined replication timing
$old_chrom = 0;
while($line = <RT>){
  @split = split ",", $line;
  if ($split[0] !~ $old_chrom and $split[0] !~ /23|24/){ # Autosomes only
    $chromosome = "chr" . $split[0];
    $seq = $inx->fetch("$chromosome");
    print $split[0], "\n";
    $old_chrom = $split[0];
  }
  if ($split[0] !~ /23|24/){
    $stop = $split[2] + 1; # To capture the nt before the start (because this is tri-nucleotide)
    $start =  $split[1] - 1; # To capture the nt after the end
    $code_seq = $seq->subseq("$start", "$stop");
    @sequence = split "", $code_seq;

    for($a = 0; $a < scalar @sequence - 2; $a++){
	$tri = $sequence[$a] . $sequence[$a+1] . $sequence[$a+2];
	if($tri !~ /N/ and $sequence[$a+1] =~ /C|T/ and $sequence[$a] =~ /A|C|T|G/ and $sequence[$a+2] =~ /A|C|T|G/){
	  $tri_context_of_RT_genome{$tri}++;
	}
	if($tri !~ /N/ and $sequence[$a+1] =~ /A|G/ and $sequence[$a] =~ /A|C|T|G/ and $sequence[$a+2] =~ /A|C|T|G/){
	  $tri = $sequence[$a+2] . $sequence[$a+1] . $sequence[$a]; #invert it
	  $tri =~ tr/ATGC/TACG/; #complement it
          $tri_context_of_RT_genome{$tri}++;
        }
    }
  }
}
close RT;

# Fetching reference genome nucleotides
# Finding WG tri distribution for normalization
for($b = 1; $b <= 22; $b++){
  $chr = "chr" . $b;
  if($b == 23){$chr = "chrX";}
  if($b == 24){$chr = "chrY";}
  $seq = $inx->fetch("$chr");
  $start = 1;
  $stop = ($seq->length()) - 1; # In case we hit the end of the chromosome
  $code_seq = $seq->subseq("$start", "$stop");
  @sequence = split "", $code_seq;
 
  for($a = 0; $a < scalar @sequence - 2; $a++){
    $tri = $sequence[$a] . $sequence[$a+1] . $sequence[$a+2];
    if($tri !~ /N/ and $sequence[$a+1] =~ /C|T/ and $sequence[$a] =~ /A|C|T|G/ and $sequence[$a+2] =~ /A|C|T|G/){
      $tri_context_of_ref_genome{$tri}++;
    }
    if($tri !~ /N/ and $sequence[$a+1] =~ /A|G/ and $sequence[$a] =~ /A|C|T|G/ and $sequence[$a+2] =~ /A|C|T|G/){
      $tri = $sequence[$a+2] . $sequence[$a+1] . $sequence[$a]; #invert it
      $tri =~ tr/ATGC/TACG/; #completment it
      $tri_context_of_ref_genome{$tri}++;
    }
  }
}

# Get denominator (number of nt essentially as RT genome is pruned
foreach $tri (keys %tri_context_of_RT_genome){
  $correction_1 = $correction_1 + $tri_context_of_RT_genome{$tri};
  $correction_2 = $correction_2 + $tri_context_of_ref_genome{$tri};
}
close OUT;

# Calcuate the correction bias. These are proportions of relative trinucleotide frequencies
open(OUT, ">correction_values.csv");
foreach $tri (keys %tri_context_of_RT_genome){
  print OUT $tri, ",";
  $correction = ($tri_context_of_RT_genome{$tri}/$correction_1) / ($tri_context_of_ref_genome{$tri}/$correction_2);
  print OUT $correction, "\n";
  $correction_values{$tri} = $correction;
}
close OUT;
undef(%tri_context_of_RT_genome);
undef(%tri_context_of_ref_genome);

# Signatures are those for a cell type. For Bcell, this SBS 1,5,9, and 40. 
open(SIG, "/path/to/Bcell_reduced_V3.2_COSMIC_signatures.tsv");
open(OUT, ">corrected_WG_Bcell_reduced_V3.2_COSMIC_signatures.tsv");
while($line = <SIG>){
  chomp($line);
  @split = split "\t", $line;
  if($line =~ /Type/){ print OUT $line, "\n";}
  if($line !~ /Type/){ 
    print OUT $split[0], "\t", $split[1], "\t", $split[2], "\t";
    print OUT $split[3] * $correction_values{$split[1]}, "\t";
    print OUT $split[4] * $correction_values{$split[1]}, "\t";
    print OUT $split[5] * $correction_values{$split[1]}, "\t";
    print OUT $split[6] * $correction_values{$split[1]}, "\n";
  }
}
close OUT;
close SIG;

# There is another step afterwards where corrected signatures are normalized so that the sum of 
# trinucleotide frequencies adds up to 1. 
```
