## Supplementary material for "Multifactorial heterogeneity of the human mutation landscape related to DNA replication dynamics": fit_signatures_whole_genome.R

```
#######################################################
# Caballero et al. 2022
# Description: Calculates trinucleotides
#  and fits signatures
#######################################################

# Note, fitting signatures by replication timing bin follows the
# same steps performed in each bin.

## INPUTS:
INPUT_MUTATIONS = read.csv(file= "/path/to/mutations/csv", header = FALSE)
INPUT_RT = read.csv(file = "/path/to/replication/timing/csv", header = FALSE)

# LIB:
'%ni%' <- Negate('%in%')
library(OneR)
library(data.table)
library(mltools)
library(binr)
library(ggpubr)
library(MASS)
library(NMF)
library(MutationalPatterns)


## Step 1: Get trinucleotide counts

INPUT_MUTATIONS$V8[INPUT_MUTATIONS$V1 == "chrX"] = NA
INPUT_MUTATIONS$V8[INPUT_MUTATIONS$V1 == "chrY"] = NA

types = table(as.character(INPUT_MUTATIONS$V11)) 				
mutation_types = row.names(types)						
cat("Number of mutation types: ", length(mutation_types), "\n") # Check to be 96
output.matrix = matrix(nrow = length(mutation_types), ncol = 2)
output.matrix[,1] = mutation_types

for (snp in 1:length(mutation_types)){ output.matrix[snp,2] = types[[snp]]}
write.table(output.matrix[,2], file = "mutation_matrix.csv", sep = ",", row.names = output.matrix[,1])


## Step 2: NMF transform counts into signatures

SIGNATURES <- read.delim(file = "/path/to/corrected_WG_Bcell_reduced_V3.2_COSMIC_signatures.norm.tsv", header = TRUE)
MUTATION_MATRIX = read.csv(file = "mutation_matrix.csv", header = TRUE, row.names = 1)

MUTATION_MATRIX = MUTATION_MATRIX + 0.001 
new_order = match(row.names(MUTATION_MATRIX), SIGNATURES$Somatic.Mutation.Type)
SIGNATURES = SIGNATURES[as.vector(new_order),]
row.names(SIGNATURES) = SIGNATURES$Somatic.Mutation.Type
SIGNATURES = as.matrix(SIGNATURES[,4:length(SIGNATURES)])

FIT_SIG <- fit_to_signatures(MUTATION_MATRIX, SIGNATURES)
contribution = as.matrix(FIT_SIG$contribution)

# Run to get the cosine sim
# cos_sim_matrix(MUTATION_MATRIX, FIT_SIG$reconstructed) 

# Bootstrap statistic of fit
bootstrap_statistic <- fit_to_signatures_bootstrapped(MUTATION_MATRIX, SIGNATURES, n_boots = 1000, method = "regular")

# Turn counts into proportions
bootstrap_statistic = as.matrix(bootstrap_statistic) / sum(MUTATION_MATRIX)
FIT_SIG$contribution = FIT_SIG$contribution / sum(MUTATION_MATRIX)

max_bount_bootstrap = vector()
min_bount_bootstrap = vector()
Standard_dev_boot = vector()
for (j in 1:ncol(bootstrap_statistic)){Standard_dev_boot = c(Standard_dev_boot, sd(bootstrap_statistic[,j]))}

max_bount_bootstrap = c(max_bount_bootstrap, Standard_dev_boot)
min_bount_bootstrap = c(min_bount_bootstrap, Standard_dev_boot)

df = data.frame(x= row.names(FIT_SIG$contribution), y = as.vector(FIT_SIG$contribution), mi = min_bount_bootstrap, ma = max_bount_bootstrap)
df$x = factor(df$x, levels = c("SBS1", "SBS5", "SBS9", "SBS40"), ordered = TRUE)

# Print result:
df
```
