## Supplementary material for "Multifactorial heterogeneity of the human mutation landscape related to DNA replication dynamics": mutation_counts_vs_binned_RT.R

```
#######################################################
# Caballero et al. 2022
# Description: Taking described bin limits and
#  partitioning mutations into those bins. 
#######################################################

## INPUTS:
args = commandArgs(trailingOnly=TRUE)
bin_number = as.numeric(args[1])  # "20"
name = as.character(args[2])  # "LCL"
mut_file_loc = as.character(args[3])  # Location of mutation csv
rt_file_loc = as.character(args[4])  # Location of replication timing csv

## LIB:
'%ni%' <- Negate('%in%')
library(OneR)
library(data.table)
library(mltools)
library(binr)
options(stringsAsFactors = FALSE) 

INPUT_MUTATIONS_LCL = read.csv(file= mut_file_loc, header = FALSE)
INPUT_RT = read.csv(file = rt_file_loc, header = FALSE)

chromosomes = c("chr1", "chr2","chr3", "chr4", "chr5", "chr6", "chr7", "chr8", 
	"chr9", "chr10", "chr11", "chr12", "chr13", "chr14", "chr15", "chr16", 
	"chr17", "chr18", "chr19", "chr20", "chr21", "chr22");

nsamp = length(unique(as.character(INPUT_MUTATIONS_LCL$V7)))

# Autosomes only
INPUT_MUTATIONS_LCL$V8[INPUT_MUTATIONS_LCL$V1 == "chrX"] = NA;
INPUT_MUTATIONS_LCL$V8[INPUT_MUTATIONS_LCL$V1 == "chrY"] = NA;
INPUT_RT$V5[INPUT_RT$V1 >= 23] = NA;

bins_red = vector();
x = bins(na.omit(INPUT_RT$V5), target.bins=bin_number, minpts = 10)
x$breaks <- bins.getvals(x)
write.table(data.frame(table(x$breaks)), file = "bins_out.txt")

# This command takes the "bins_out.txt" file and formats it into a simple row-based txt file
 system("perl ~/format.breaks.pl bins_out.txt > temp_bin.txt")

tab = x$binct                                                      
input = read.csv(file = "temp_bin.txt", header = FALSE)
bins = c(-4, sort(input$V1), 4)                           
m = as.numeric(INPUT_MUTATIONS_LCL$V8[!is.na(INPUT_MUTATIONS_LCL$V8)])
y = cut(m, breaks= bins)                             
z=table(y)
mut = vector();
for(i in 1:nrow(z)){ mut = c(mut, z[[i]]); }  # Vectorizing the bin counts
bins_red = vector()
for(i in 2:length(bins)){
  get_mean = mean(as.numeric(INPUT_RT$V5[((as.numeric(INPUT_RT$V5) > bins[i-1]) & (as.numeric(INPUT_RT$V5) <= bins[i]))]), na.rm = TRUE)
  cat(name, ",", get_mean, ",", mut[i-1], "\n", sep = "")
}

# Mutation counts per bin are captured as > output
```
