## Supplementary material for "Multifactorial heterogeneity of the human mutation landscape related to DNA replication dynamics": mutation_density_continuous_across_chromosome.R

```
#######################################################
# Caballero et al. 2022
# Description: Plotting mean number of mutations
#  in a sliding window across the chromosome.
#######################################################

## INPUTS:
bin_size = 1000000
step = 500000

args = commandArgs(trailingOnly=TRUE)
MUTATIONS = read.csv(file= args[1], header = FALSE)  # Mutations csv
RT = read.csv(file = args[2], header = FALSE)  # Replication timing csv

## LIB :
library(ggplot2)
'%ni%' <- Negate('%in%')
library("OneR")
library(data.table)
library(mltools)
library('binr')


chromosomes = c("chr1", "chr2","chr3", "chr4", "chr5", "chr6", "chr7", "chr8",
        "chr9", "chr10", "chr11", "chr12", "chr13", "chr14", "chr15", "chr16",
        "chr17", "chr18", "chr19", "chr20", "chr21", "chr22");

## Step 1: Getting mean number of mutations across the chromosome in defined bins+step
for (Chr in 1:22){
  rt = (RT$V1 == Chr)                               
  coor = as.numeric(RT$V2[rt])
  interval = ceiling((max(coor) - min(coor))/step);

  vec_count = vector();
  vec_coor = vector();
  for (i in 0:(interval - 1)){
     min_val =  min(coor) + (i*step)
     max_val =  min(coor) + (i*step) + bin_size

     mid_point = (max_val + min_val) / 2

     samples = MUTATIONS$V7[(MUTATIONS$V2[MUTATIONS$V1 == chromosomes[Chr]] > min_val) & 
	(MUTATIONS$V2[MUTATIONS$V1 == chromosomes[Chr]] <= max_val)];
     t <- c(table(as.character(samples)), rep(0, (length(unique(MUTATIONS$V7)) - length(table(as.character(samples))))))
     cat(Chr, ",", mid_point, ",", mean(t), "\n", sep = "")
  }
}


## Step 2: Calculate correlation to replication timing
rt_values = vector();
mut_values = vector();
for(Chr in 1:22){
  rt = (RT$V1 == Chr) & !is.nan(RT$V5);
  rt_df <- data.frame(x = RT$V2[rt], y = RT$V5[rt]);
  mut = (MUTATIONS$V1 == Chr)
  mut_df <- data.frame(x = MUTATIONS$V2[mut], y =  MUTATIONS$V3[mut]);

  # This inverts and normalizes data to the scale of RT
  mut_df$y[(mut_df$y == 0)] = NaN;
  mut_df$y = mut_df$y  - mean(mut_df$y, na.rm = TRUE)
  mut_df$y = mut_df$y / sd(mut_df$y, na.rm = TRUE)
  mut_df$y = mut_df$y * -1

  mut_values = c(mut_values, mut_df$y);
  for (win in 1:length(mut_df$x)){
    A = which.min(abs(rt_df$x - mut_df$x[win]));
    rt_values = c(rt_values, rt_df$y[A])
  }
  sub = !is.nan(mut_values)
  C = cor(mut_values[sub], rt_values[sub])
}

cat(args[1], ",", args[2], ",", C, "\n", sep = "")
```
