## Supplementary material for "Multifactorial heterogeneity of the human mutation landscape related to DNA replication dynamics": phase_mutations_from_bam.pl

```
#######################################################
# Caballero et al. 2022
# Description: Phasing a bam and extracting mutations.
#   This one is formatted to work for the iHART
#   data which is on AWS.
#
# This script is designed to work for an individual file
# for parallel operations.
#######################################################
#!usr/bin/perl

# INFILES:
$MUTATIONS = "/path/to/mutations/csv/";
# Note, for iHART, mutations are lifted to hg19 as these BAM
# files are aligned to hg19. 

$ID = $ARGV[0]; # Such as NA12878
$BAM = $ARGV[1]; # Path to sample BAM, this can be an s3:// loc on AWS (samtools is okay with that)
$POSITIONS = $ID . "_positions.tsv";

# OUTFILES:
$out = "$ID.phased.mutations.csv";
open(OUT, ">$out");
$mut_out = "$ID.mut.pileup";
$pos_out = "$ID.pos.pileup";


open(MUT, $MUTATIONS);
while($mutationline = <MUT>){
  chomp($mutationline);
  @mutsplit = split ",", $mutationline;
  @info = split "_", $mutsplit[6];
  $mut_id = $info[scalar @info - 1];
    # For a mutation of this sample
    if($mut_id =~ $ID){
      $mutsplit[0] =~ s/chr//g;
      # Getting the read IDs that cover the exact coordinate of the mutation from the BAM.
      # The mpileup command will tell us the allele and the sourced read name. 
      $string = "samtools mpileup --output-QNAME -a -r $mutsplit[0]\:" . $mutsplit[1] . "\-" . $mutsplit[1] . " " . 
       $BAM . "\> " . $mut_out;
      system($string);

      open(POS, $POSITIONS);  # These are mat/pat phased SNPs
      while($posline = <POS>){	
	@possplit = split "\t", $posline;
        if($possplit[0] =~ /^$mutsplit[0]$/){ # Matching chromosome
	  if(abs($possplit[1] - $mutsplit[1]) <= 1000){ # SNPs within 1000bp of mutation (very generous, could just be 2x read length)
	    $loc = $possplit[0] . "_" . $possplit[1];
            $posstore{$loc}++;
            # Identifying which 
            if($possplit[3] =~ /M/){$mat_allele{$loc} = $possplit[2];}
            if($possplit[3] =~ /P/){$pat_allele{$loc} = $possplit[2];}
            # The file has two lines per location (a Mat and Pat line with the repective allele. Only performing pileup command
            # once both are recorded, hence the == 2.
            if($posstore{$loc} == 2){
              # Again, grabbing the location of the phased SNPs, alleles, and respective read name.
  	      $string = "samtools mpileup --output-QNAME -a -r $possplit[0]\:" . $possplit[1] . "\-" . $possplit[1] . " " . 
	        $BAM . "\> " . $pos_out;
              system($string);

              # Parsing the phased SNP pileup and sorting reads with maternal allele and paternal allele. 
              open(POS_PILEUP, $pos_out);
	      while($pileupline = <POS_PILEUP>){
	        chomp($pileupline);
                @inpileup = split "\t", $pileupline;
                $inpileup[4] =~ s/\$//g;
                $inpileup[4] =~ s/\^.//g;

                @alleles = split "", $inpileup[4];
 	        @RG = split ",", $inpileup[6];
	        for($a = 0; $a < scalar @alleles; $a++){
                  if(uc($alleles[$a]) =~ uc($mat_allele{$loc})){
                    $read_id{$RG[$a]} = "MATERNAL";
		  }
                  if(uc($alleles[$a]) =~ uc($pat_allele{$loc})){
                    $read_id{$RG[$a]} = "PATERNAL";
                  }
                }
              }
              close POS_PILEUP;
            }
	  }
       }
     }
     close POS;
  
    # Now that we know read names with maternal and paternal ID, we compare to the reads collected before
    # that contain the mutation allele.
    open(POS, $mut_out);
    while($pileupline = <POS>){
      chomp($pileupline);
      @inpileup = split "\t", $pileupline;
      $inpileup[4] =~ s/\$//g;
      $inpileup[4] =~ s/\^.//g;

      @alleles = split "", $inpileup[4];
      @RG = split ",", $inpileup[6];
      $maternal = 0;
      $paternal = 0;
      for($a = 0; $a < scalar @alleles; $a++){
        if(uc($alleles[$a]) =~ uc($mutsplit[4])){ # We are at a mutation allele
          if($read_id{$RG[$a]} =~ /PATERNAL/){ # Mut allele is on a paternal read
            $paternal = 1;
          }
          if($read_id{$RG[$a]} =~ /MATERNAL/){ # Mut allele is on a maternal read
            $maternal = 1;
          }
        }
      }
    }
    close POS; 

    # If exclusive to one parent, phased. If found on both, ambiguous. Of on neither, unknown. 
    if($paternal == 1 and $maternal == 0){ print OUT $mutationline, "\t", "PATERNAL", "\n";}
    if($paternal == 0 and $maternal == 1){ print OUT $mutationline, "\t", "MATERNAL", "\n";}
    if($paternal == 0 and $maternal == 0){ print OUT $mutationline, "\t", "UNKNOWN", "\n";}
    if($paternal == 1 and $maternal == 1){ print OUT $mutationline, "\t", "AMBIG", "\n";}
    undef(%read_id);
  }
}
close MUT;
close OUT;
system("rm $POSITIONS");
system("rm $mut_out");
system("rm $pos_out");
```
