## Supplementary material for "Multifactorial heterogeneity of the human mutation landscape related to DNA replication dynamics": README

This is a small collection of the code for Caballero et al. 2022. These files represent the most critical and complex code. 

The full project uses a mix of Perl, R, MATLAB, and Python. All scripts have a descriptive header. 

Other scripts used in this project are available upon request. 
