## Supplementary material for "Multifactorial heterogeneity of the human mutation landscape related to DNA replication dynamics": simulate_clustering_by_replication_timing.R

```
#######################################################
# Caballero et al. 2022
# Description: Simulating expected clustering
#  based on the number of mutations in RT bins. 
#  Performed 1000 iterations.
#######################################################

## INPUTS:
RANGE <- read.csv(file = "SHM_mut_counts.csv", header = FALSE);
# This file contains the replication timing range and the number
# of SHM context mutations (last column). 
# SHM,-4,-1.744592,-1.969652,26913

## LIB:
library(ClusteredMutations)
library(seriation)

for (r in 1:length(RANGE$V1)){
  system(paste0("rm sim/", r, ".sim.clusters.csv")) # Clearing an old file since this uses append concat
  for (sim in 1:1000){
    system(paste0("shuf -n ", RANGE$V5[r], " motifs/", r, ".SHM.motifs.csv > rand.csv"));
    mut_in_clusters = 0
    num_clusters = 0
    num_mut = 0

    RAND_COOR = read.csv(file = "rand.csv", header = FALSE)
    for(chr in 1:22){
	coordinates = RAND_COOR$V2[RAND_COOR$V1 == chr]	
	num_mut = num_mut + length(coordinates)
	clusters <- showers(data = NULL, chr = NULL, position = as.numeric(coordinates), min = 2, max = 500)
	mut_in_clusters = mut_in_clusters + sum(clusters$number)
	num_clusters = num_clusters + length(clusters$number)			
    }
    cat(sim, ",", mut_in_clusters, ",", num_clusters, "\n", file = paste0("sim/", r, ".sim.clusters.csv"), sep = "", append = TRUE)
  }
}
```
