## Supplementary material for "Multifactorial heterogeneity of the human mutation landscape related to DNA replication dynamics": strand_bias_ratio_in_replication_timing_bins.R

```
#######################################################
# Caballero et al. 2022
# Description: Calculates strand bias ratios
#  for mutation pathways in LCL
#######################################################

## INPUTS:
LEAD <- read.csv(file = "contributions.lead.txt", header = TRUE)
LAG <- read.csv(file = "contributions.lag.txt", header = TRUE)

## Single signatures
signatures = unique(LEAD$Signature)
replication_times = unique(LEAD$x) # Bins
for (sig in signatures){
  subsig = LEAD$Signature == sig
  ratios = log2(LEAD$y[subsig]/LAG$y[subsig])
  a = LEAD$y[subsig]/LAG$y[subsig]
  rel_error = a* sqrt((LEAD$ma[subsig] / LEAD$y[subsig])^2 + (LAG$ma[subsig]/ LAG$y[subsig])^2)
  error =  log2(a) - log2(a - rel_error)

  df = data.frame(x = replication_times, y = ratios, mi = ratios-error, ma = ratios+error)
  print(df)
}


# Signatures combined into pathway (e.g. clock-like and SHM)
Signature = vector()
x = vector()
y = vector()
ma = vector()
mi = vector()

for (t in replication_times){
  sub_Clock = ((LEAD$Signature == "SBS1") | (LEAD$Signature == "SBS5") | (LEAD$Signature == "SBS40") ) & (LEAD$x == t)
  Clock_mut_lead = sum(LEAD$y[sub_Clock])
  Clock_mut_lag = sum(LAG$y[sub_Clock])
  Clock_ratio = log2(Clock_mut_lead / Clock_mut_lag)

  a = Clock_mut_lead / Clock_mut_lag
  Clock_lead_error = sqrt(sum(LEAD$ma[sub_Clock]^2)) #SEM
  Clock_lag_error = sqrt(sum(LEAD$ma[sub_Clock]^2)) #SEM
  rel_error = a* sqrt((Clock_lead_error / Clock_mut_lead)^2 + (Clock_lag_error/ Clock_mut_lag)^2)
  Clock_error =  log2(a) - log2(a - rel_error)

  sub_SHM = (LEAD$Signature == "SBS9")  & (LEAD$x == t)
  SHM_mut_lead = sum(LEAD$y[sub_SHM])
  SHM_mut_lag = sum(LAG$y[sub_SHM])
  SHM_ratio = log2(SHM_mut_lead / SHM_mut_lag)

  a = SHM_mut_lead / SHM_mut_lag
  SHM_lead_error = sqrt(sum(LEAD$ma[sub_SHM]^2)) #SEM
  SHM_lag_error = sqrt(sum(LEAD$ma[sub_SHM]^2)) #SEM
  rel_error = a* sqrt((SHM_lead_error / SHM_mut_lead)^2 + (SHM_lag_error/ SHM_mut_lag)^2)
  SHM_error =  log2(a) - log2(a - rel_error)

  Signature = c(Signature, "Clock", "SHM")

  y = c(y, Clock_ratio, SHM_ratio)
  ma = c(ma, Clock_ratio+Clock_error, SHM_ratio+SHM_error)
  mi = c(mi, Clock_ratio-Clock_error, SHM_ratio-SHM_error)
  x = c(x, t, t)
}

df = data.frame(x =x, y = y, mi = mi, ma = ma, sig = Signature)
df$sig= factor(df$sig, levels = c("Clock", "SHM"), ordered = TRUE)
print(df)
```
