## Supplementary material for "Multifactorial heterogeneity of the human mutation landscape related to DNA replication dynamics": TIGER_note_hg19to38_liftover.m

```
%%%%%%%%%%%%%%%%%%%%%%%%%%%%%%%%%%%%%%%%%%%%%%%%%%%%%%%
% Caballero et al. 2022
% Description: Modified TIGER resource generation
%  to note which coordinates in hg38 do not have
%  an hg19 equivalent on the same chromosome. 
%%%%%%%%%%%%%%%%%%%%%%%%%%%%%%%%%%%%%%%%%%%%%%%%%%%%%%%

clear;clc

% VARIABLES:
genome_build='hg38';
read_length=100;
win_size=2500;

resources = '../tiger_resources/scripts/';
addpath(resources); % The special scripts like seg filt are here too
resources = '..//tiger_resources/';
addpath(resources);

load hg38_chr_length
load hg38_sequence

disp('Loading filtered reads')
for Chr=1:length(chr_lengths)
    disp(['Chromosome ' num2str(Chr)])
    filename=cell2mat(['Chr' chrnum(Chr) '_' num2str(read_length) 'bp_filt.txt']);
    fid=fopen(filename);
    C=textscan(fid,'%f');  % read coordinate
    fclose('all'); clear fid filename

    % all genomic coordinates that are not uniquely alignable
    Coordinates_to_remove_align=setdiff(1:chr_lengths(Chr),C{1}); 

    % all genomic coordinates that failed an hg19 to 38 liftover
    filename = cell2mat(['/fs/cbsukoren2/storage/mc2698/9_Individual_RT/resource_liftover/chr' chrnum(Chr) '_failed_lifted_coordinates.txt']);
    fid=fopen(filename);
    C=textscan(fid,'%f');  % read coordinate
    fclose('all'); clear fid filename

    Coordinates_to_remove{Chr,1}(:,1) = sort(unique([cell2mat(C); transpose(Coordinates_to_remove_align)]));
    clear C Coordinates_to_remove_align
end

eval(['mkdir ' resources 'Alignability_and_GC_filters/lift_' genome_build '_' num2str(read_length) 'bp']) 
eval(['cd ' resources 'Alignability_and_GC_filters/lift_' genome_build '_' num2str(read_length) 'bp'])
eval(['save -v7.3 Coordinates_to_remove_' genome_build '_' num2str(read_length) 'bp Coordinates_to_remove'])

% load file with chromosome lengths
load Gap_hg38.mat Gap 

disp('Calculating windows')
for Chr=1:length(chr_lengths)
    disp(['Chromosome ' num2str(Chr)])
    L2=setdiff(1:chr_lengths(Chr),Coordinates_to_remove{Chr});
    Wins{Chr,1}(:,2)=L2(1:win_size:end-win_size+1); % start 
    Wins{Chr,1}(:,3)=L2(win_size+1:win_size:end)-1; % end
    Wins{Chr,1}(:,1)=Chr;
    Wins{Chr,1}(:,4)=round(mean(Wins{Chr}(:,2:3)')'); % window center
    
    % remove windows that span gaps
    G=Gap(Gap(:,1)==Chr,2:3);
    in=[];
    for i=1:size(G,1)
        in=[in;find(Wins{Chr,1}(:,2)<=G(i,1) & Wins{Chr,1}(:,3)>=G(i,2))];
    end
    Wins{Chr}(in,:)=[];
    
end

eval(['save ' genome_build '_' num2str(read_length) 'bp_wins_'   num2str(win_size)  ' Wins'])

quit()
```
