## Supplementary material for "Multifactorial heterogeneity of the human mutation landscape related to DNA replication dynamics": README

Caballero et al. 2022. 

Column descriptions for each file:
1. Chromosome. LS180 and HT115 only go up to 22. CLL, HCT116, and LCL go up to X (which is coded as 23). 
2. Start coordinate of replication timing window. 
3. End coordinate of replication timing window. Each window is 2.5kb of uniquely alignable sequence. 
4. Center coordinate of replication timing window. 
5. Mean replication timing value for all samples.
6. Replication timing value for all male samples (not included in LS180, HT115, and HCT116).
7. Replication timing value for all female samples (not included in LS180, HT115, and HCT116).

All replication timing values are plotted on a Z-score with a mean of 0 and a standard deviation of 1.
All coordinates are in hg38.
